# Injectable Nanoparticle-Based Hydrogels Enable the Safe and Effective Deployment of Immunostimulatory CD40 Agonist Antibodies

**DOI:** 10.1101/2021.06.27.449987

**Authors:** Santiago Correa, Emily C. Gale, Aaron T. Mayer, Zunyu Xiao, Celine Liong, John H. Klich, Ryanne A. Brown, Emily L. Meany, Olivia M. Saouaf, Caitlin L. Maikawa, Abigail K. Grosskopf, Joseph L. Mann, Juliana Idoyaga, Eric A. Appel

## Abstract

When properly deployed, the immune system can eliminate deadly pathogens, eradicate metastatic cancers, and provide long-lasting protection from diverse diseases. Unfortunately, realizing these remarkable capabilities is inherently risky as disruption to immune homeostasis can elicit dangerous complications or autoimmune disorders. While current research is continuously expanding the arsenal of potent immunotherapeutics, there is a technological gap when it comes to controlling when, where, and how long these drugs act on the body. Here, we explore the ability of a slow-releasing injectable hydrogel depot to reduce the problematic dose-limiting toxicities of immunostimulatory CD40 agonist (CD40a) while maintaining their potent anti-cancer efficacy. We leverage a previously described polymer-nanoparticle (PNP) hydrogel system that exhibits shear-thinning and yield-stress properties that we hypothesized would improve locoregional delivery of the CD40a immunotherapy. Using PET imaging, we demonstrate that prolonged hydrogel-based delivery redistributes CD40a exposure to the tumor and the tumor draining lymph node (TdLN), thereby reducing weight loss, hepatotoxicity, and cytokine storm associated with standard treatment. Moreover, CD40a-loaded hydrogels mediate improved local cytokine induction in the TdLN and improve treatment efficacy in the B16F10 melanoma model. PNP hydrogels, therefore, represent a facile, drug-agnostic method to ameliorate immune-related adverse effects and explore locoregional delivery of immunostimulatory drugs.

## 1. Introduction

Although cancer immunotherapy has led to remarkable outcomes for certain patients,^[1]^ the majority do not respond to current immunotherapy strategies.^[2]^ Many of the non-responders exhibit poorly immunogenic “cold” tumors, which feature numerous and often non-redundant mechanisms of immunosuppression.^[3]^ These suppressive adaptations work together to prevent effective T cell-mediated anti-tumor immunity.^[4]^ For these patients to benefit from immunotherapy, alternative strategies are required to sensitize their tumors to the immune system.^[5]^

Immunostimulatory drugs (*e.g.*, CD40 agonists, TLR agonists, cytokines, bispecific antibodies) may provide a path towards transforming cold tumors into an immunogenic “hot” phenotype.^[6]^ These drugs broadly activate the innate immune system and overwhelm immunosuppressive mechanisms at play in the tumor microenvironment (TME) and tumor draining lymph nodes (TdLNs).^[2b, 3b]^ Notably, leveraging immunostimulants appears to be critical to provoking anti-tumor immune responses in a variety of malignancies.^[7]^ Yet, these drugs also activate potent immune responses throughout normal and healthy tissues, causing serious and dose-limiting immune-related adverse effects (IRAEs).^[1b, 8]^ Consequently, immunostimulation is a double-edged sword in immuno-oncology, which creates an outstanding need for technological solutions to safely deploy these drugs as part of combination therapies.

CD40 agonist antibodies (CD40a) are particularly promising immunostimulants which have been explored in clinical trials. CD40a therapy works through a variety of mechanisms that deplete immunosuppressive myeloid cells, repolarize tumor-associated macrophages (TAMs), and improve T cell priming by antigen presenting cells (APCs).^[9]^ Taken together, CD40a therapy can both expand the pool of anticancer T cells and transform the tumor microenvironment into a more hospitable landscape for those T cells to carry out their functions **(Scheme 1)**. Moreover, CD40a may prove to be a critical component for effective combination immunotherapy, where it has shown promising early data in clinical trials.^[10]^ Unfortunately, concerns remain with the safety and general tolerability of CD40a,^[11]^ as with immunostimulants in general. In particular, cytokine storms, infusion reactions, low platelet counts, neutropenia and liver toxicity have been carefully monitored in CD40a clinical trials.^[10a, 12]^ The abundance of endogenous off-target CD40 also creates a considerable antigen sink effect that contributes to poor overall pharmacokinetics.^[13]^

**Scheme 1.**
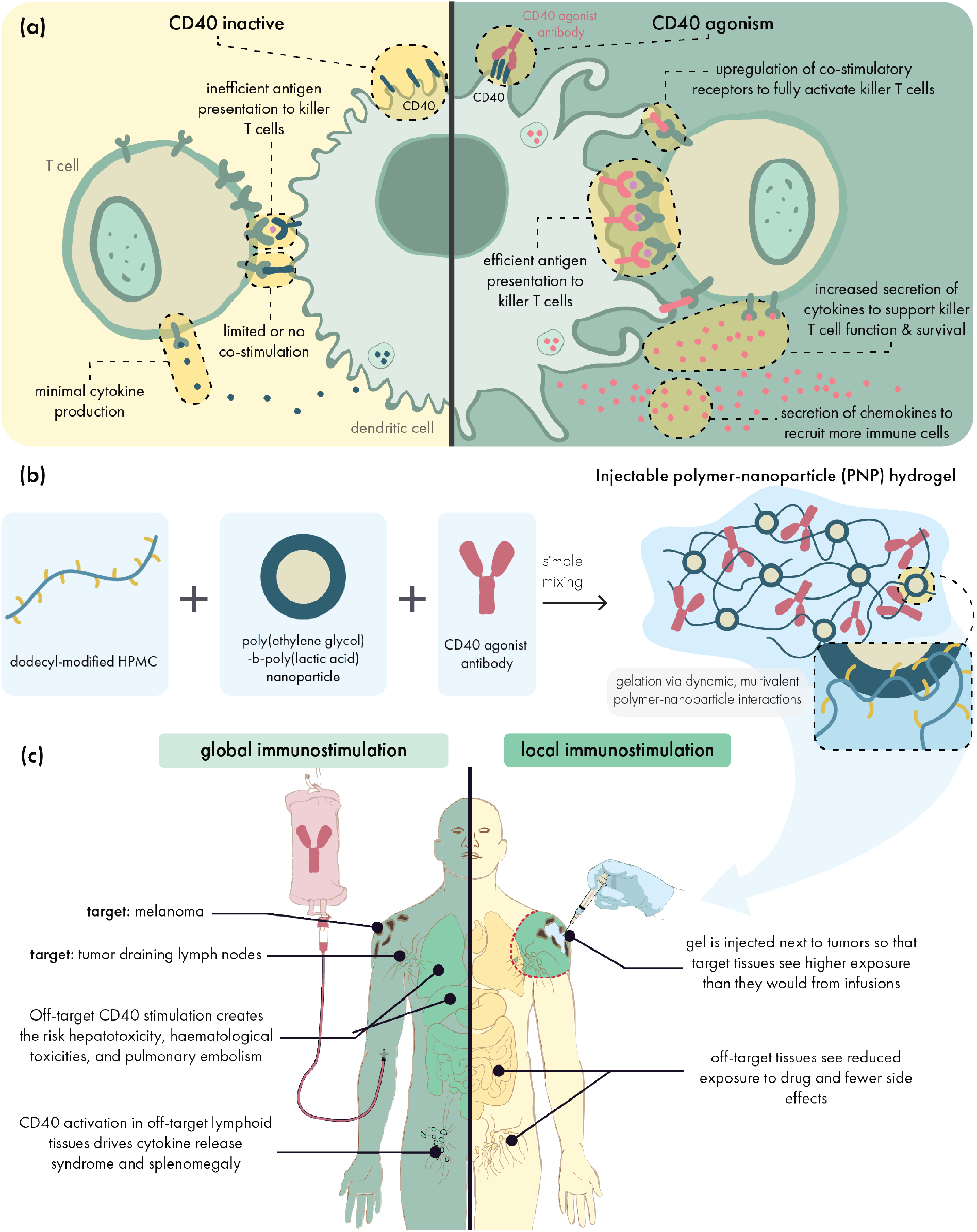
Hydrogels can localize the effects of CD40 agonists to reap their anti-tumor benefits while abating their immune-related adverse effects. **(A)** CD40 agonist antibodies engage CD40 receptors on antigen presenting cells to potently upregulate antigen presentation, co-stimulatory receptor expression, and secretion of immunostimulatory cytokines. Altogether, these effects potentiate T cell priming and create a more supportive environment for effector T cell function. **(B)** Injectable, supramolecular polymer-nanoparticle (PNP) hydrogels composed of dodecyl-modified hydroxypropyl methylcellulose and poly(ethylene glycol)-*b*-poly(lactic acid) nanoparticles can be used to encapsulate CD40 agonists for local drug delivery. **(C)** Peritumoral hydrogel administration leads to slow release of CD40 agonists into the local microenvironment, focusing the immunostimulatory effects on the tumor and draining lymphatics. In contrast, traditional systemic approaches lead to widespread exposure throughout the body and the occurrence of immune-related adverse effects.

Here, we investigated how the sustained locoregional delivery of CD40a could facilitate safer and more effective outcomes compared to conventional administration methods. Based on our prior results with sustained and local vaccine delivery,^[14]^ we hypothesized that injectable polymer-nanoparticle (PNP) hydrogels could also provide a highly localized and extended release of CD40a. We evaluated this approach in the poorly immunogenic B16F10 melanoma model, which exhibits an immune desert phenotype that fails to respond to frontline PD-1/L1 checkpoint inhibitor immunotherapy.^[15]^ By quantitatively tracking biodistribution using positron emission tomography (PET), we found that hydrogels substantively altered antibody pharmacokinetics to favor drug exposure in target tissues. Locoregional hydrogel delivery circumvented serious limitations facing CD40a antibodies, including poor pharmacokinetics due to antigen sink effects as well as dose limiting IRAEs that include acute weight loss, cytokine storm, and hepatotoxicity. In addition to safety benefits, locoregional CD40a therapy increased drug exposure to the TME and TdLNs, thus generating a more potent immune response in the relevant tissues compared to systemic administration. Correspondingly, hydrogel-mediated locoregional CD40a therapy was more tolerable and effective even than local bolus injections and synergized with PD-L1 blockade. Notably, PNP hydrogels improved safety and efficacy without requiring modification of CD40a, and thus they may be broadly beneficial for the delivery of other immunostimulants.

## 2. Results

### 2.1. Preparation of immunostimulatory polymer-nanoparticle (PNP) hydrogels

We hypothesized that the sustained, local delivery of CD40a from injectable hydrogels would increase drug exposure in the tumor and its draining lymph nodes (Scheme 1). Prolonged stimulation of antigen-presenting cells (APCs) in these target tissues was expected to maximize the efficacy of CD40a therapy. Meanwhile, reduced systemic exposure was expected to reduce toxicity. To this end, we prepared CD40a-loaded PNP hydrogels by physically mixing aqueous solutions of dodecyl-modified hydroxypropyl methylcellulose (HPMC-C_12_), poly(ethylene glycol)- block-poly(lactic acid) (PEG-b-PLA) nanoparticles (NPs), and CD40a (Scheme 1b).

Dynamic, multivalent interactions between HPMC-C_12_ and PEG-b-PLA NPs generate a crosslinked supramolecular hydrogel exhibiting robust solid-like properties (G’ greater than G’’) across a range of physiologically relevant frequencies (**Figure 1a**). We recently demonstrated that these nanoparticle-derived physical networks form through entropy-driven non-covalent interactions between the nanoparticle and polymer constituents, which have the benefit of providing temperature invariant mechanical properties.^[16]^ We also performed flow rheology measurements to confirm that these hydrogels exhibit shear-thinning behavior, becoming *ca.* 100-fold less viscous under high shear rate conditions (Figure 1b). This extreme shear thinning behavior is necessary for injection of the hydrogel through fine gauge needles used in clinical settings.^[17]^ After injection, the hydrogel network must self-heal to regain its solid-like properties to serve as a prolonged drug depot. To demonstrate the self-healing capabilities of this material, we measured the viscosity of the system while alternating between high and low shear rates that simulate the mechanical forces exerted during manual injection from a syringe (Figure 1c). Over multiple high-shear cycles, we observed that our hydrogels rapidly regain their solid-like properties within 2 seconds following cessation of high-shear conditions. This quick reestablishment of the hydrogel network is expected to minimize burst release following administration and permit our hydrogel to serve as a prolonged drug depot after administration.

**Figure 1.**
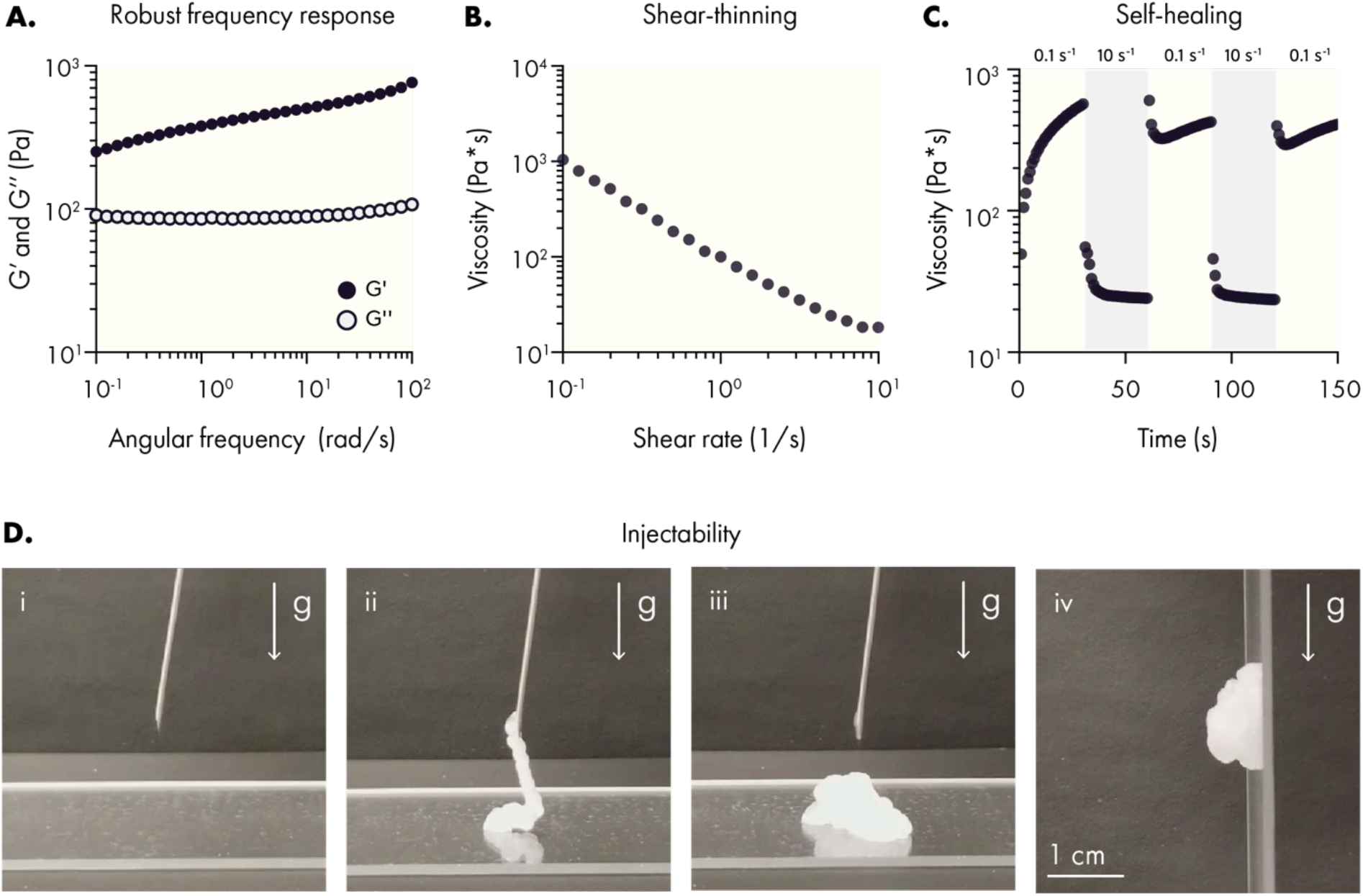
PNP hydrogels exhibit solid-like rheological properties with robust shear-thinning and self-healing capabilities. (A) Oscillatory shear rheology of PNP hydrogels indicating a storage modulus (G’) greater than the loss modulus (G’’) throughout the tested frequencies. (B) Steady shear rheology measurements indicating a decrease of viscosity at increasing shear. (C) Step-shear rheological behavior after two cycles of high shear (10 s^−1^) with intermediate low shear (0.1 s^−1^) cycles. (D) Injection of PNP hydrogel through a 21-gauge needle: (i) pre-injection, (ii) injection of the hydrogel through the needle demonstrating liquid-like rheological properties, (iii) when no longer under high shear conditions PNP hydrogels regain solid-like properties, such as (iv) the resistance of flow due to gravity.

Overall, these rheological properties indicated the antibody-loaded hydrogel became more liquid-like under high shear, allowing it to be injected through standard needle geometries for minimally invasive introduction of the drug depot. Post-injection, the material quickly recovers its solid-like mechanical properties. To further validate this behavior, CD40a PNP hydrogels were injected from a 1 mL syringe equipped with 21-gauge needle (Figure 1d), demonstrating the hydrogel is readily injected and re-solidifies into a solid-like material capable of resisting flow under gravity. Batch-to-batch characterization of these materials indicate that these rheological properties are consistent between independent batches (**Figures S1 & S2**).

### 2.2. Sustained locoregional CD40a delivery is effective against local and distant tumors

For local immunotherapy to be clinically relevant for metastatic disease, it must not only inhibit local tumors but also distant tumors. To evaluate if our CD40a hydrogels could inhibit the growth distant tumors, we explored efficacy in a syngeneic dual-flank model of B16F10 melanoma (**Figure 2a**). One tumor on each mouse was peritumorally treated with an injectable hydrogel containing either anti-PD-1 (aPD-1), CD40a, or no drug. To compare results to a clinically relevant standard, we also evaluated response to repeated systemic administration of aPD-1 antibody. We observed that a single administration of hydrogel loaded with a high dose of CD40a (100 μg, which is 20x the preclinical/clinical maximum tolerated dose (MTD) for systemic administration)^[11c]^ was able to markedly slow the progression of both B16 tumors (Figure 2b). Consistent with slowed tumor growth, mice treated with CD40a hydrogels saw a modest but significant increase in overall survival, compared to either local or systemic aPD-1 treatments. Assessment of tumor growth inhibition on day 10 indicated that CD40a hydrogel therapy had a significant therapeutic benefit for both treated and untreated tumors (Figure 2d), indicating an abscopal anti-cancer effect with CD40a hydrogel therapy. To date, the efficacy of CD40a monotherapy has been poor in clinical trials, which may be attributable to dose-limiting IRAEs of CD40a in the context of systemic administration.^[18]^ These data suggest that CD40a delivered locally may improve the clinical performance of CD40a monotherapy.

**Figure 2.**
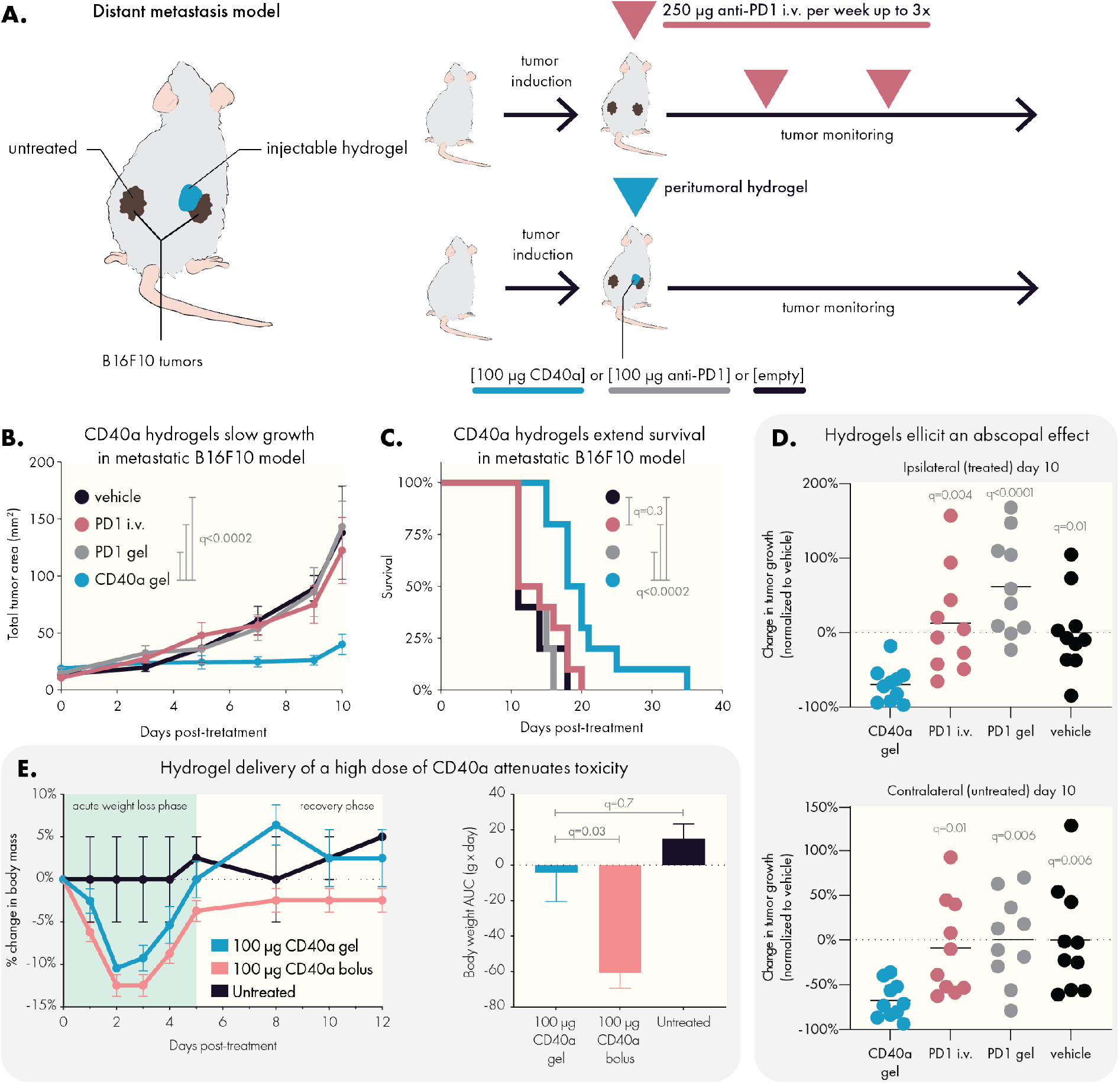
Sustained locoregional hydrogel-based delivery of CD40a slows tumor growth and improves safety in a metastatic model of B16F10 melanoma. (A) C57B/6 mice were inoculated with two flank B16F10 tumors and treated with an injectable hydrogel containing either anti-PD1 or CD40a, or were treated with weekly systemic administration of anti-PD1. Empty hydrogels (vehicle) were used as a negative control. Local delivery of a high dose (100 μg) of CD40a (B) inhibited tumor growth and (C) extended overall survival of mice. (D) Mice treated with CD40a hydrogels exhibited reduced tumor growth in both treated and untreated tumors 10 days after treatment. (E) Comparison of treatment-induced weight loss due to high dose local CD40a administered as either a local bolus or as a PNP hydrogel. Longitudinal percent change in body mass (left panel) and the corresponding area under the curve (AUC) analysis (right panel). Efficacy (n = 10) and toxicity (n = 4 for CD40a treated mice, n = 2 for untreated) datasets represent two independent experiments. Data represented by means and SEM. Differences in tumor growth and survival were assessed using general linear models with SAS statistical software. Comparison of body weights and tumor growth inhibition was performed using one-way ANOVA. False discovery rate (FDR) was controlled using the Benjamini and Hochberg method.

Given the high dose of CD40a used in this experiment, tolerability and safety of the therapy were a concern. To evaluate gross toxicity due to treatment, we measured changes in body weight over time after administration of 100 μg CD40a delivered peritumorally in either an injectable PNP hydrogel or as a bolus (Figure 2e). At this high dose, both delivery approaches exhibit an acute weight loss phase that begins within 24 hours of treatment; however, hydrogel delivery attenuates the overall weight loss seen in this acute phase. In addition to lessening the acute symptoms, hydrogel-treated mice regained their original weight within 5 days and afterwards began to gain weight (gains consistent with growth of mice of 7-8 weeks age). In contrast, mice treated with a local bolus of CD40a failed to fully recover their original body weight throughout the duration of the study.

### 2.3. PNP hydrogels redistribute CD40 agonist exposure to target tissues

Our initial observations of anti-cancer efficacy and improved tolerability were consistent with our hypothesis that our hydrogels mediate important changes in CD40a pharmacokinetics. To further validate this hypothesis, we used positron emission tomography (PET) to track radiolabeled CD40a in B16F10 tumor-bearing mice. Because antibodies persist for extended periods in the body, we labeled CD40a with the long-lived ^89^Zr isotope, which has a half-life of 3.27 days, to track its pharmacokinetics (PK) over the course of 12 days. Mice were treated with 100 μg of CD40a, delivered peritumorally via bolus injection or in a PNP hydrogel depot (**Figure 3a**).

**Figure 3.**
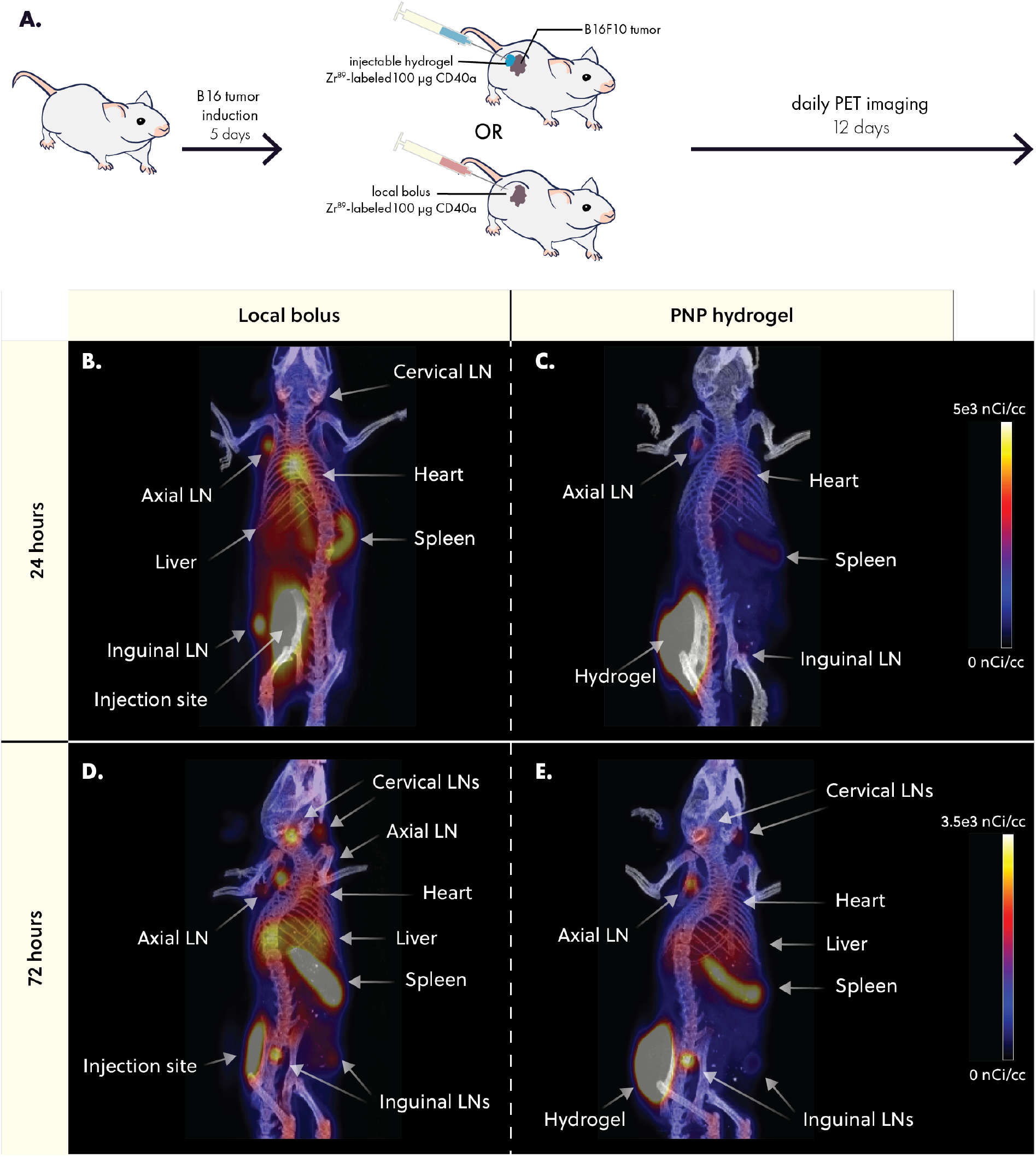
PNP hydrogels retain CD40a at the injection site and reduce exposure to distant tissues. (A) B16F10 bearing mice were treated with 100 μg of Zr^89^-labeled CD40a administered either as a PNP hydrogel or local bolus. Mice were imaged daily using positron emission tomography (PET) for 12 days. Representative images of CD40a biodistribution 24 hours after treatment for (B) local bolus or (C) PNP hydrogel administration routes. Representative images of CD40a biodistribution 72 hours after treatment for (D) local bolus or (E) PNP hydrogel administration routes. Scale bar is shared between (B) and (C), and between (D) and (E).

PET imaging revealed that hydrogel delivery led to prolonged retention of CD40a in the injection site, while bolus administration led to rapid systemic leakage and accumulation in off-target organs within 24 hours (Figure 3b-c). At 72 hours post-treatment, PNP hydrogels continued to exhibit reduced accumulation of CD40a in distant, off-target organs like the liver and spleen, compared to the local bolus (Figure 3d-e). These data show that bolus administered CD40a leaks into the systemic blood compartment before rapidly accumulating in secondary lymphoid organs throughout the body. Unlike most antibody therapies, which persist in circulation with half-lives on the order of three weeks, CD40a experiences pronounced antigen sink effects due to high levels of CD40 expressed in the spleen and lymphatics.^[13]^ In contrast, administration of CD40a in PNP hydrogel depot slowed down systemic leakage and reduced accumulation in off-target antigen sinks over the course of the study (**Figure S3**).

To quantitatively assess how drug exposure in specific tissues is altered by hydrogel delivery, we calculated the area under the drug-time curve (AUC) in each tissue over the course of the study. We then took the ratio of the AUC from hydrogel and bolus treated mice to calculate the percent change in drug exposure due to hydrogel delivery (**Figure 4a**). PET measurements were validated by gamma counter measurements of explanted tissues at the end of the study (**Figure S4**). Quantitative analysis revealed that hydrogels significantly increased CD40a exposure in target tissues. More specifically, hydrogels increased exposure at the injection site by 96 ± 13% (q = 4 x 10^−6^) and at the tumor draining lymph node by 35 ± 19% (q = 5 x 10^−3^) over the course of this 12-day study. Notably, hydrogel delivery of CD40a reduced drug exposure in all other tissues, especially in antigen-sink organs like the spleen where drug exposure was reduced by 23 ± 2% (q = 4 x 10^−6^). We also observed significant reduction in drug exposure in all off-target lymph nodes (LN), with the exception of the ipsilateral axial LN where we observed a moderate decrease that was not significant (q = 0.1). We attributed this to the ipsilateral axial lymph node’s proximity to the injection site, which likely leads to partial drainage of lymph from the hydrogel depot. Overall, these data strongly indicate that hydrogel delivery has a profound effect on CD40a pharmacokinetics, and significantly redistributes drug exposure to target tissue and away from off-target tissue, even compared to local bolus administration.

**Figure 4.**
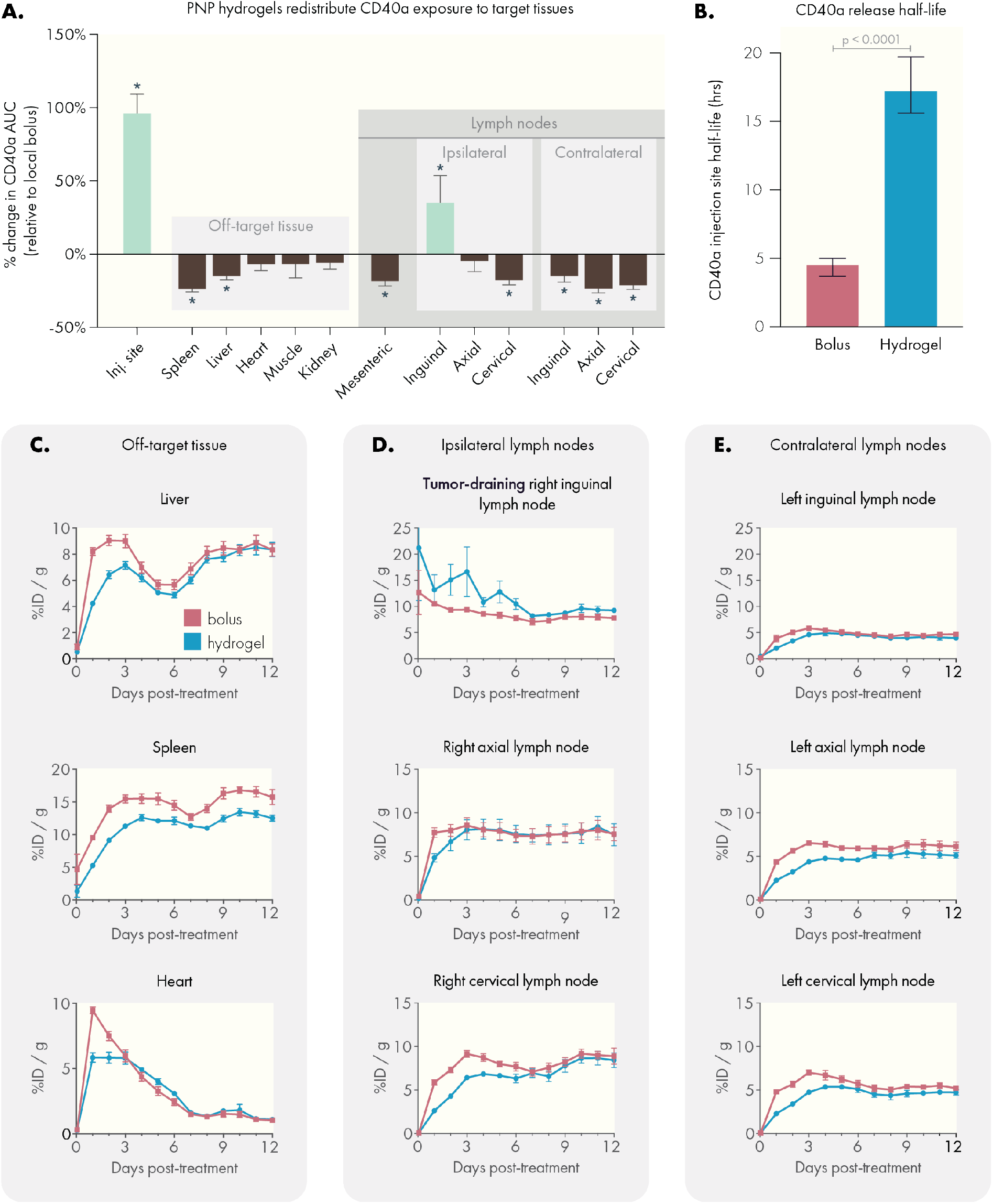
PNP hydrogels increase drug exposure to target tissues and reduce exposure in off-target tissues compared to local bolus. (A) Percent change in area under the curve (AUC) values for each tissue relative to local bolus administration. AUCs were derived from the PET pharmacokinetic data. Statistical comparisons indicate a significant difference from local bolus AUC, performed as multiple unpaired t-tests. Corrections for multiple comparisons were performed using the FDR approach (Q=1%). (B) CD40a retention half-life at the injection site, estimated from a one-phase decay fit of PET data. Comparison performed using the extra sum-of-squares F test. PET pharmacokinetic curves tracking CD40a concentration in (C) off-target tissues, (D) ipsilateral lymph nodes (including the tumor-draining lymph node), and (E) off-target contralateral lymph nodes. N=4 for both groups, data shown as mean and SEM.

Tissue-specific pharmacokinetic curves provided further insight into the way PNP hydrogels shift the biodistribution of CD40a. At the injection site, local bolus administration led to rapid CD40a leakage, with 90% of the drug released by 48 hours (**Figure S5**). In contrast, hydrogel delivery led to a sustained release of CD40a, reaching 90% release by five days. Fitting these data with a single-phase exponential decay model estimates a *ca.* 4-fold increase (p = 0.0001) in CD40a half-life at the injection site (Figure 4b), with hydrogels exhibiting a half-life of 17.2 hrs (95% CI 15.6-19.7 hrs) compared to a half-life of 4.5 hrs (95% CI 3.7-5 hrs) for local bolus administration.

Additionally, hydrogel delivery significantly reduced peak CD40a concentrations in the spleen (22.3% C_max_ reduction, q = 0.005) and heart (35.9% C_max_ reduction, q = 0.002; Figure 4c, **Figure S6**). Notably, drug concentration in the heart, which is indicative of circulating CD40a, demonstrates that circulating antibody levels are strongly depressed when delivered via hydrogel. Consistent with the antigen-sink effects documented for CD40a, circulating antibody levels rapidly decay. Highly perfused organs like the liver and spleen exhibit two peaks during the study, where the initial peak is potentially due to contribution from circulating antibody. The subsequent peak for the liver and spleen is likely due to accumulation of antibody on CD40 expressing cells which are enriched in both organs. Elimination of CD40a via the liver also contributes to the rise in drug in this tissue from 6-12 days. However, in the initial 6 days following administration, we observed a 22.1% reduction in C_max_ (q = 0.012) which may provide important benefits for mitigating the hepatotoxicity associated with this therapy.

In the ipsilateral lymph nodes (those on the same side as the injection site), we observed a rapid increase and plateau of CD40a levels in the cervical, axial, and tumor-draining inguinal lymph nodes when administering the antibody as a local bolus (Figure 3d). In general, ipsilateral LNs plateaued between 5 and 10% initial dose (ID)/gram of tissue. Following bolus administration, the tumor draining LN briefly sustains levels greater than 10% ID/g during the initial 24 hours, later decaying to ∼8% ID/g. In contrast, hydrogel delivery of CD40a sustains elevated CD40a levels (on average greater than 10% ID/g) in the tumor-draining lymph node over 6 days. We attribute the higher C_max_ and delayed decay to the slow release of CD40a from the hydrogel, which allows for turnover of CD40 in the TdLN to regenerate new binding sites for the antibody. In contrast, these data suggest that CD40a introduced via bolus administration quickly saturates local binding sites, leaving a large unbound fraction of antibody that is likely quickly shuttled back into systemic circulation via lymphatic drainage.

Consistent with our hypothesis that CD40a accumulation into distant lymph organs occurs after locally administered drug re-enters systemic circulation, we observed a delay in C_max_ in lymph nodes when CD40a was delivered via hydrogel (Figure S6D). For the ipsilateral axial and cervical LNs, which do not efficiently drain the injection site, C_max_ was reached on days 4.5 and 8.5, respectively, following local bolus administration. In contrast, hydrogel administration delayed C_max_ in these tissues to days 8 and 10.75, respectively. Drug exposure was further reduced for the contralateral LNs (those on the opposite side from the injection site). In these tissues, CD40a exposure was significantly reduced using hydrogel delivery, with C_max_ reductions of *ca.* 77% in the contralateral inguinal LN and *ca.* 35% in the contralateral axial and cervical LNs, relative to the C_max_ of their ipsilateral counterparts. In terms of kinetics, bolus administration led to the rapid uptick of CD40a in contralateral LNs by 24 hours, while hydrogel delivery led to a slow increase in drug concentration over 3-4 days. Altogether, these PK experiments suggest that some CD40a reaches distant ipsilateral LNs via drainage, accounting for increased drug exposure in these tissues; however, the primary driver of accumulation is likely leakage into systemic circulation, explaining why bolus administration quickly reaches plateaus in these tissues while hydrogel delivery delays plateaus in all distant lymph tissues.

### 2.4. PNP hydrogels attenuate toxic side effects associated with CD40a

Although we observed safety benefits at a high dosage of CD40a, we conducted additional studies to identify a minimally toxic hydrogel dosage. Based on our PET study, it was apparent that the local tissues, and in particular the tumor draining lymph node, have a finite capacity to absorb and retain CD40a. With this in mind, we hypothesized that a dose titration would identify a dose which would be retained efficiently in the target tissues, preventing the spill-over we observed at later time points with the 100 μg dose and maximizing the safety of the treatment. We assessed several indicators of gross toxicity including treatment-induced weight loss and histopathology of the liver and spleen. Using two syngeneic cancer models (MC38 and B16F10), we tracked body weight changes following CD40a treatment administered as either a local bolus, systemic bolus, or hydrogel. Locoregional therapy was evaluated at three CD40a doses (50, 25, and 10 μg), while systemic therapy was limited to the MTD of 5 μg. For these toxicology studies, we focused on weight loss occurring in the acute toxicity phase noted in our prior high-dose study (1 to 5 days post-treatment), as even the 100 μg dose hydrogels mediated healthy weight gain after the acute toxicity period. We observed that hydrogel-based CD40a delivery significantly reduced acute weight loss compared to dose-matched local bolus controls (**Figure 5A, Figure S7**). While some degree of weight loss was noted in all treatments, we observed that a 10 μg dose of CD40a delivered in a hydrogel depot induced minimal weight loss that is not statistically different from isotype controls or the systemic MTD. In contrast, local bolus administration of 10 μg CD40a still caused significant weight loss compared to isotype controls (q = 0.016).

**Figure 5.**
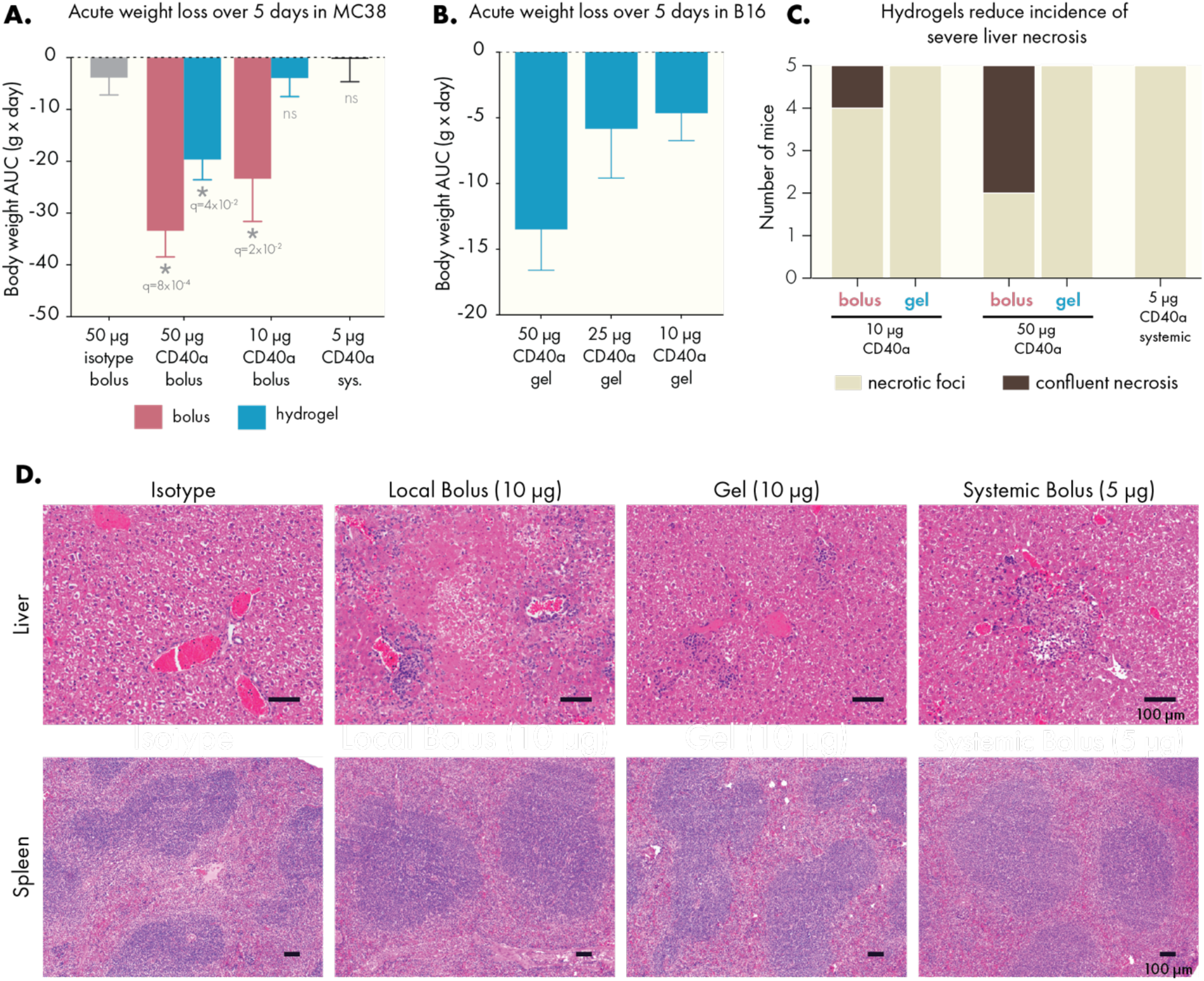
PNP hydrogels mitigate gross toxicity caused by CD40a therapy. CD40a dose was titrated and treatment-induced weight loss was observed. Area under the curve analysis of body mass curves during the acute weight loss phase (5 days post-treatment) for mice bearing (A) MC38 and (B) B16F10 tumors. Statistical comparisons performed with a one-way ANOVA, multiple testing error controlled using the FDR approach (Q = 5%). Histopathology of the liver and spleen were assessed in B16F10 mice, 3 days after treatment. (C) Quantification of degree of liver necrosis. (D) Representative images of livers and spleens from treated mice. Scale bars indicate 100 microns. Results are from 3 independent experiments. N = 8-10 for (A), n = 4-5 for (B), and n = 5 for (C-E).

CD40a therapy has been shown to induce severe hepatotoxic side effects,^[11a]^ and it can also lead to enlargement of the spleen and other lymphoid organs. To assess how hydrogel delivery impacted toxicity in these tissues, mice were sacrificed 3 days after treatment and their tissues were weighed, fixed, and analyzed by a pathologist (Figure 5C-E, **Figure S8**). Livers from all mice treated with CD40a exhibited necrotic foci, defined as regions smaller than 0.4 mm in area. Yet, only mice treated with bolus administration of CD40a exhibited more serious confluent necrosis (areas of necrosis > 0.4 mm). All mice treated with CD40a exhibited expansion of the splenic white pulp, and spleen and TdLN enlargement correlated strongly to CD40a dose (**Figure S9 and S10**). Indicators of toxicity were reduced greatly by reducing the overall dose of CD40a, and the effect of dose reduction on toxicity were stronger when administering the antibody in a hydrogel depot. Importantly, inhibition of tumor growth and overall survival were not impacted by reducing the dose of CD40a, indicating dose-sparing benefits to local therapy (**Figure S11**). Overall, these data indicate that local hydrogel delivery allows the CD40a dose to be safely increased 2-fold relative to the systemic maximum tolerated dose.

### 2.5. Hydrogel delivery of CD40a mitigates treatment-induced cytokine storm

Cytokine storm syndrome is a common side effect of immunostimulant therapy and manifests shortly after CD40a administration.^[8b]^ To assess how hydrogel delivery may alter the induction of systemic cytokines, we collected blood from mice 24 hours after treatment and analyzed serum using a Luminex 48-plexed cytokine assay (**Figure 6**, **Figure S12**). This time point was chosen due to the rapid onset of cytokine storm observed in CD40a clinical trials.^[10a, 12]^ We evaluated dose-response using either a local bolus or hydrogel administration route and again compared results to systemic administration of the MTD.

**Figure 6.**
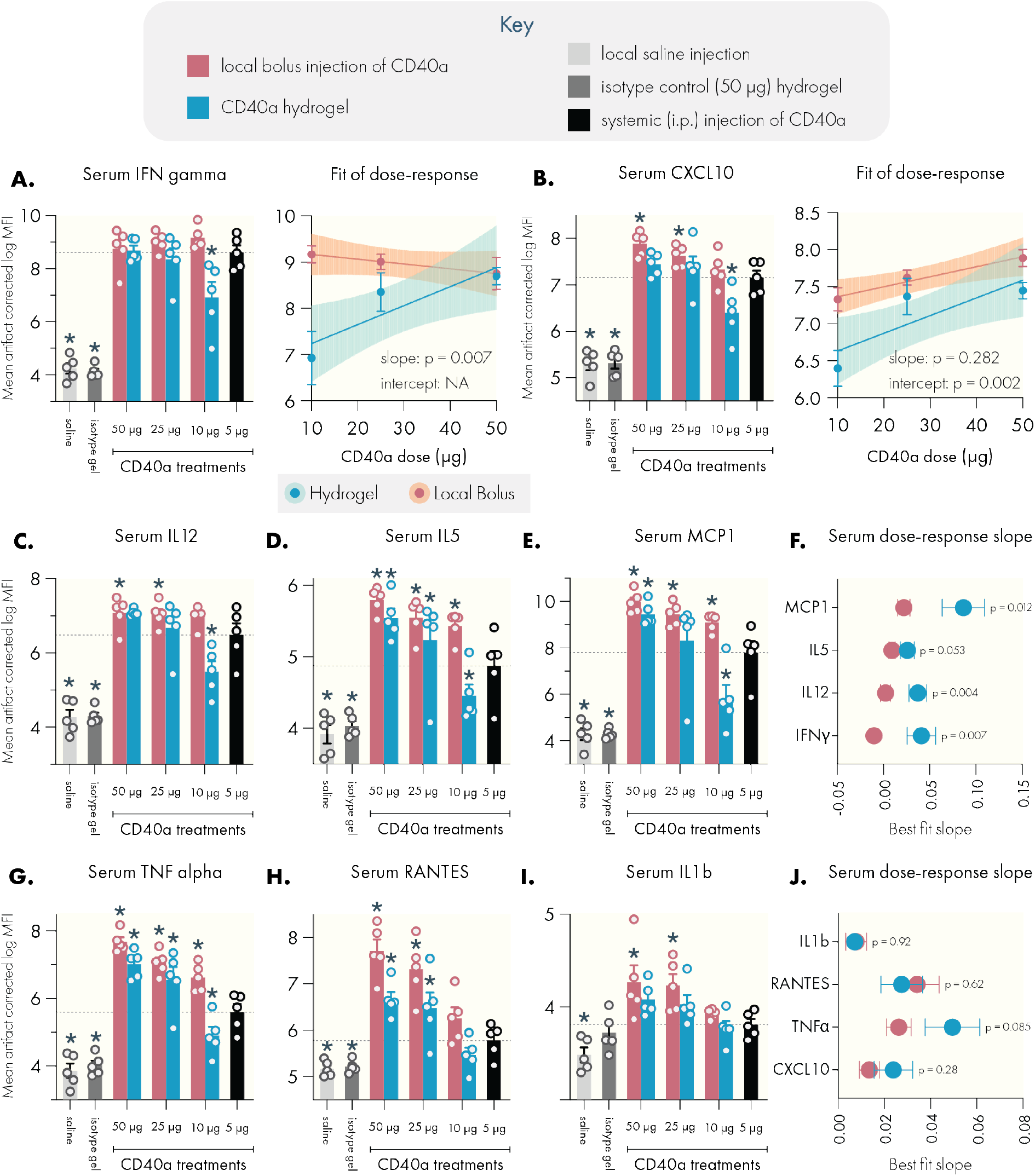
PNP hydrogels attenuate the induction of pro-inflammatory serum cytokines compared to dose-matched local bolus administration. Serum cytokine levels were assessed by Luminex 24 hours after treatment. Systemic levels of cytokine-storm associated cytokines (A) IFNγ, with linear regression of dose-response data indicating a significant change in the dose-response slope associated with administration route. (B) CXCL10, with linear regression of dose-response data indicating a significant change in the elevation of the dose-response curve associated with administration route. Hydrogel delivery suppressed systemic induction of (C) IL12, (D) IL5, and (E) MCP1. (F) Linear regression revealed significant differences between administration methods in the dose-response slope for these cytokines. Hydrogel delivery also suppressed systemic induction of (G) TNFα, (H) RANTES, and (I) IL1b. (J) The dose-response slope is not altered in these cytokines, however the elevation of the dose-response curve is significantly altered depending on administration route (see Figure S15). Dotted lines indicate mean value corresponding to the 5 μg systemic dose (maximum tolerated systemic dose). N = 5 for all groups, data represented as mean and SEM. Shaded area in linear regressions indicate 95% CI. * Denote a significant difference (p < 0.05) from the systemic MTD. Multiple testing error was controlled using the FDR approach (Q = 5%). Statistical comparison of slope and intercept performed using built-in analysis in the linear regression functionality of GraphPad Prism.

Estimation of serum cytokine concentrations from raw Luminex mean fluorescence intensity (MFI) values indicate a profound induction of pro-inflammatory cytokines (Figure S12). Consistent with the mechanism of CD40 agonism, IFNγ levels were the most potently induced. Saline and isotype treated mice exhibited IFNγ levels below 1 pg/mL, while mice treated with the systemic MTD exhibited average IFNγ levels of 134 pg/mL. All doses administered as a local bolus elevated serum IFNγ to similar levels. However, hydrogel delivery of 10 μg CD40a in a hydrogel suppressed levels more than 6-fold to an average of 21 pg/mL. At this dose, hydrogels consistently mediated comparable or significantly depressed systemic levels of proinflammatory cytokines, including *ca.* 4-fold reductions in average systemic TNFα, IL12, and MCP1 levels.

However, interpretation of pg/mL values from Luminex data can be biased due to experimental artifacts and differences between standards and real biological samples.^[19]^ To more accurately infer true biological patterns in the data, we instead analyzed the raw fluorescence signal using a utility developed by the Stanford Human Immune Monitoring Center to correct for plate, batch, lot, and nonspecific binding artifacts in Luminex data (**Figures S13 and S14**).^[20]^ Based on these corrected values, we found that hydrogel-based CD40a delivery substantively altered the induction of systemic cytokines by altering either the slope or the elevation of the dose-response curve (**Figure 6A-B, Figure S15**). In both cases, the end result is a significant suppression of peripheral cytokine levels when administering CD40a in a hydrogel. Importantly, at the 10 μg dose, hydrogel therapy resulted in reduced or comparable systemic cytokine induction relative to the 5 μg systemic MTD control.

In particular, the slopes of the dose-response curves for IFNγ, IL12, IL22, and MCP1 were significantly steeper for hydrogel therapies relative to local bolus administration (p < 0.02), indicating that serum cytokine induction falls off more quickly for hydrogel therapies (Figure S15). The dose-response curves for TNFα, IL6, IL1b, IL5, IL18, CXCL10, RANTES, and CXCL1 indicated similar slopes, but significantly lower elevation for the hydrogel curves. To further bolster these analyses, we performed hierarchical clustering of serum cytokine levels induced by CD40a treatments, which revealed two major clusters that corresponded to low and high cytokine induction (**Figure S16**). Notably, 4/5 of 10 μg hydrogel treated mice clustered along with the low induction cluster, while none of the dose-matched local bolus treated mice fell into this category. Interestingly, 4/5 of mice receiving the 5 μg systemic control, which most closely simulates the conventional clinical approach,^[12]^ clustered into the high induction cluster. Along with our toxicological data, these results further validate that PNP hydrogels enable the safe delivery of higher doses of CD40a. In particular, administering CD40a therapy *via* hydrogel mitigated serum cytokine induction by altering dose-response dynamics, which may provide important safety benefits for CD40 agonists and other potent immunostimulants.

### 2.6. Locoregional administration improves induction of effector cytokines in tumor draining lymph nodes

Although high levels of cytokines in the periphery can exacerbate immune related adverse effects, sufficient levels of effector cytokines must be induced in the vicinity of the tumor to mount a successful immune response. To determine if hydrogel administration sufficiently drives local cytokine induction, we recovered TdLNs from mice three days after treatment and analyzed cytokine levels using the Luminex cytokine assay (**Figure 7**). The 3 day timepoint was chosen based on our PET pharmacokinetic data for the injection site and was anticipated to provide a more robust comparison between the sustained and rapid-release local delivery approaches. As with the serum cytokine analysis, we provide the results of this assay in quantitative numbers (**Figure S17**), but we rely on the more accurate analysis based on the corrected fluorescence signal for our biological inferences (**Figures S18 and S19**).

**Figure 7.**
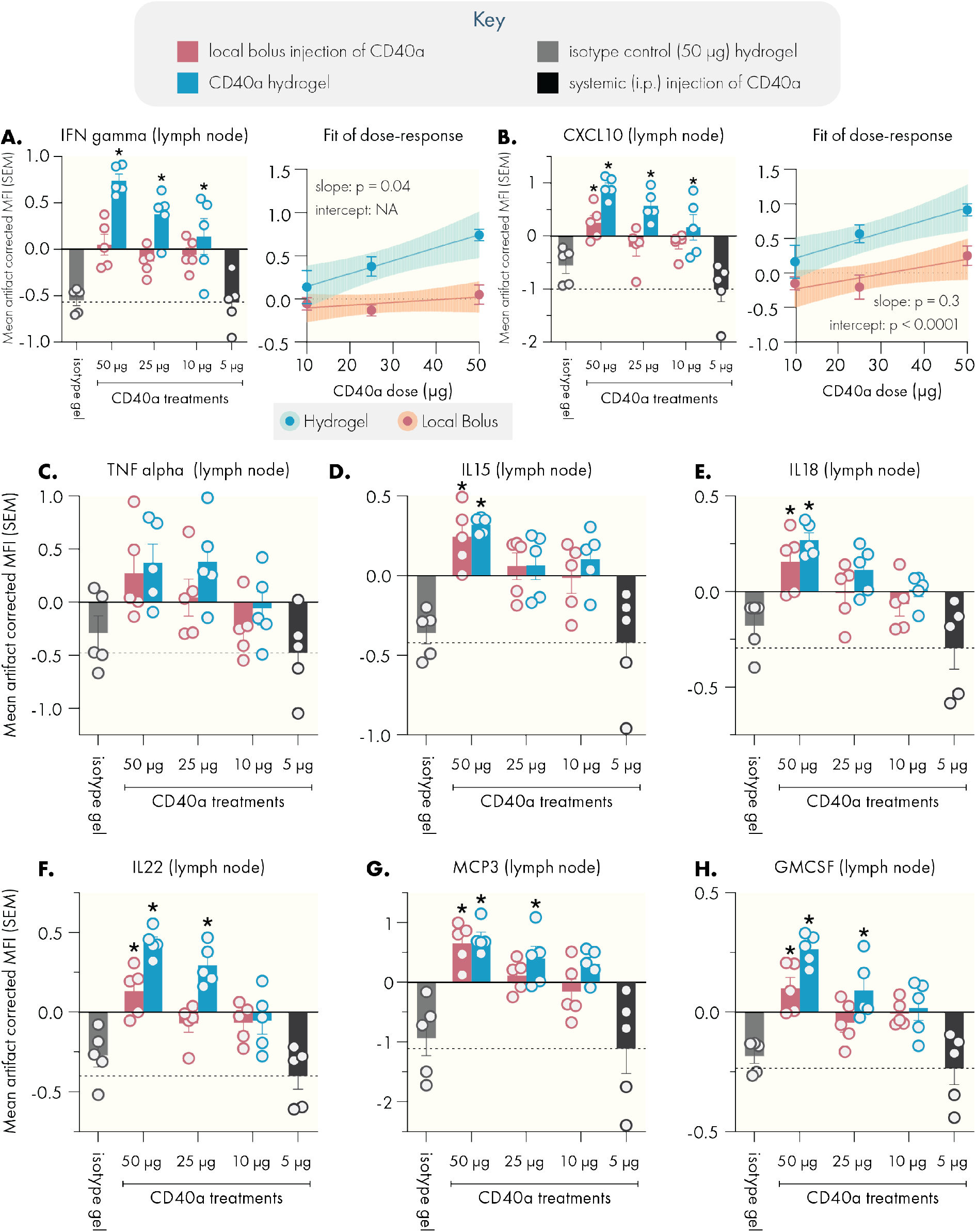
Sustained delivery of CD40a via PNP hydrogel increases effector cytokine levels in the tumor draining lymph node. Lymph nodes were collected from mice 3 days after treatment for cytokine analysis by Luminex. Hydrogel delivery of CD40a led to elevated levels of the pro-inflammatory cytokine IFNγ and the associated CXCL10 chemokine. (A) IFNγ levels with linear regression indicating a significant change in the dose-response slope associated with administration method. (B) CXCL10 levels with linear regression indicating a significant change in the elevation of the dose-response curve associated with administration method. Compared to systemic routes, locoregional delivery elevated levels of immunostimulatory cytokines (C) TNFα, (D) IL15, (E) IL18, and (F) IL22. Similarly, locoregional delivery enhanced levels of the chemokines (G) MCP3 and (H) GMCSF. Dotted line indicates mean value corresponding to the 5 μg systemic dose (maximum tolerated systemic dose). N = 5 for all groups, data represented as mean and SEM. * Denote a significant difference (p < 0.05) from the MTD systemic dose. Multiple testing error was controlled using the FDR approach (Q = 5%). Statistical comparison of slope and intercept performed using built-in analysis in the linear regression functionality of GraphPad Prism.

In contrast to the results for systemic cytokine induction, we found that hydrogels tended to drive higher levels of effector cytokines in TdLNs. Notably, both local bolus and hydrogels were able to induce greater cytokine levels compared to systemic administration of the MTD. However, we observed a consistent trend where hydrogels induced similar or improved local cytokine induction, compared to the dose-matched local bolus. We observed significantly (p < 0.05) elevated levels of the key cytokines IFNγ and CXCL10 at the 10 μg dose for hydrogels, which are critical for initiating a strong Th1 immune response (Figure 7A-B).

To further characterize how hydrogel delivery alters local cytokine induction, we performed linear regressions on the dose-response data for the cytokines most strongly induced by CD40a therapy (**Figure S20**). Compared to the serum cytokine data, hydrogel and local bolus approaches generate dose-response curves with similar slopes, except for IFNγ where the hydrogel dose-response curve exhibited a sharply steeper slope (p = 0.04). Notably, local bolus delivery exhibits an IFNγ dose-response curve that is relatively insensitive to increases in CD40a dose. In contrast, IFNγ induction in response to hydrogel delivery appeared to be quite sensitive to the encapsulated dose. We attribute this difference to the ability of the hydrogels to sustain the release of CD40a over time, which appears to increase production of IFNγ beyond what is possible with local bolus techniques. Because IFNγ is a key driver in the immunological response to CD40 agonism, these elevated levels may in turn explain the elevation noted for other cytokines. We noted significant elevation of the dose-response curve (p < 0.04) for IL18, IL22, CXCL10, MCP3, and GMCSF when CD40a was delivered in a hydrogel depot. Unlike with the serum cytokine data, hierarchical clustering of the TdLN data did not reveal interpretable clusters (**Figure S21**), which may be because fewer cytokines appeared to be induced at this timepoint. Overall, these data indicate that locoregional therapy, and especially hydrogel-mediated therapy, can significantly increase effector cytokine levels in the vicinity of the tumor, compared to conventional clinical approaches.

### 2.7. Hydrogels enable dose-sparing strategies for CD40a monotherapy

Although high-dose CD40a hydrogel monotherapy led to efficacy and reduced toxicity in the B16F10 model, our pharmacokinetic and pharmacodynamic data suggested that further safety benefits could be achieved by reducing the locoregional dose. Our initial dose-sparing studies demonstrated similar tumor growth inhibition and overall survival benefits across 100, 50, 25, and 10 μg doses when delivered via PNP hydrogels (Figure S11). Based on our toxicology and cytokine induction studies, the 10 μg dose of CD40a is also significantly safer when delivered in a hydrogel depot, compared to a local bolus. While this is the lowest dose we evaluated, this locoregional dose is 2-fold higher than the systemic preclinical/clinical MTD for CD40a drugs. To determine whether the hydrogel delivery vehicle may be necessary for maintaining efficacy at these lower doses, we evaluated treatment efficacy in the immunotherapy-resistant B16F10 model of melanoma using the 10 μg dose of CD40a delivered either in a hydrogel or as a local bolus (**Figure 8A**). We found that a single administration of 10 μg of CD40a could significantly slow down tumor growth (q < 0.005) of established B16F10 tumors (avg. diameter = 4 ± 2 mm), but only if it was administered in a hydrogel depot (Figure 8B). Consistent with tumor growth data, CD40a hydrogels, but not CD40a local bolus, were able to significantly extend median survival in this model (q = 0.009) and yielded one long-term survivor (Figure 8C). As expected, hydrogels loaded with CD40a isotype controls had no effect on tumor outgrowth. These data indicate that hydrogel delivery may provide advantages for revisiting the clinical feasibility of CD40a monotherapy.

**Figure 8.**
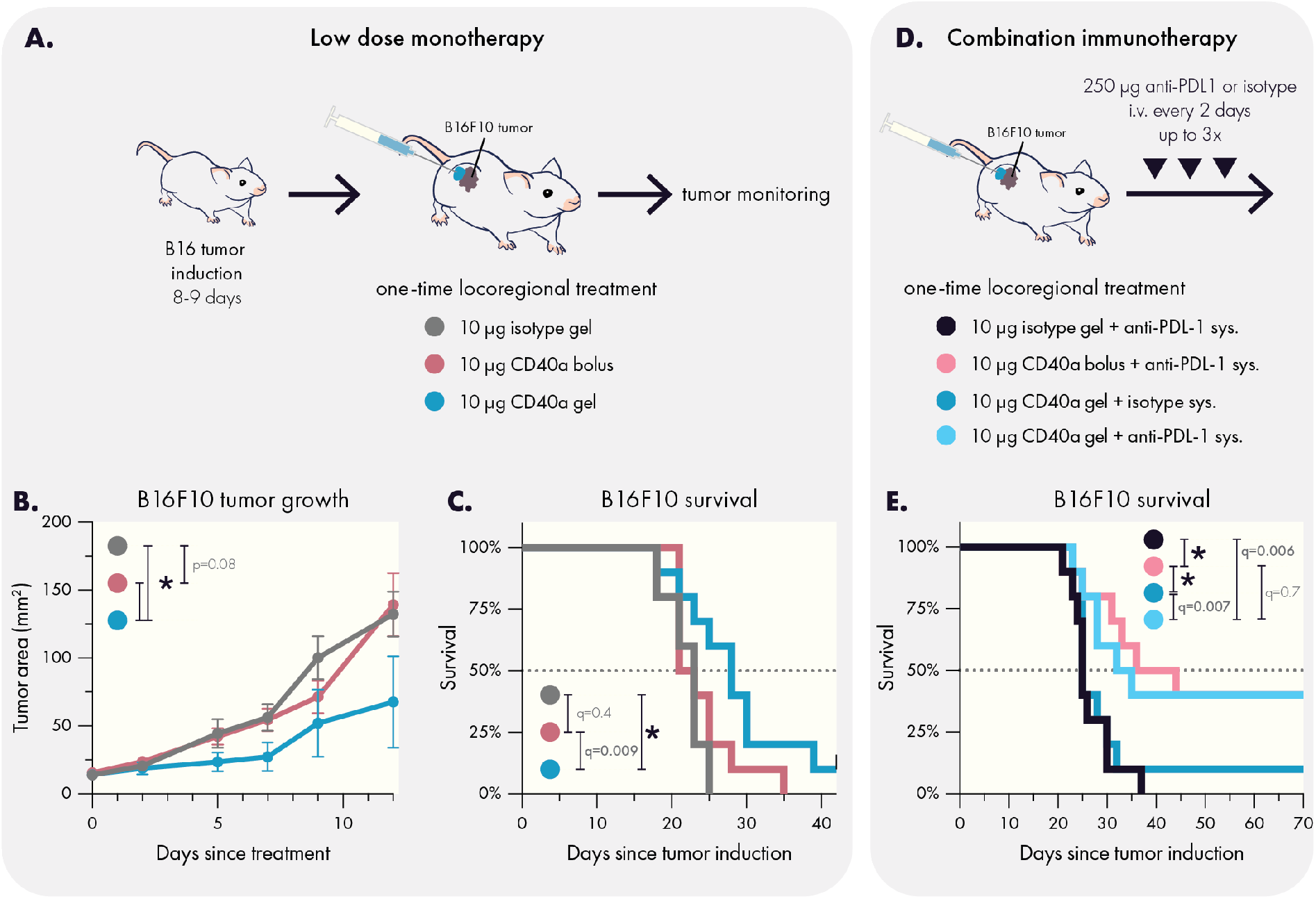
Hydrogel delivery yields effective low dose CD40a monotherapy and synergizes with PD-L1 blockade. (A) Experimental scheme for low dose monotherapy on established B16F10 tumors. Hydrogels containing 10 μg CD40a slowed the (B) growth of B16F10 tumors and (C) extended survival. (D) Experimental scheme for evaluating the impact of additional systemic anti-PD-L1 therapy in the treatment of established B16F10 tumors. (E) Low dose locoregional CD40a therapy synergizes with systemic PD-L1 blockade and yields long term survivors in the B16F10 model. Results are from two independent experiments. N=10 for all groups. Data in (B) indicates mean and SEM. * Indicates adjusted p values < 0.005. Differences in tumor growth and survival were assessed using general linear models with SAS statistical software. False discovery rate (FDR) was controlled using the Benjamini and Hochberg method (Q=1%).

### 2.8. Locoregional CD40a therapies synergize with PD-L1 blockade

There are compelling clinical data to suggest that CD40a-based combination immunotherapy may hold promise for treating historically incurable cancers.^[10a]^ In particular, treatments that combine PD-1/L1 blockade appear to be quite promising. To determine whether locoregional CD40a therapy may synergize with checkpoint blockade, we evaluated how a single administration of CD40a coupled to systemic anti-PD-L1 therapy would impact survival in the immunotherapy-resistant B16F10 model (Figure 8D). Intriguingly, we observed that both local bolus and hydrogel administration of CD40a synergized strongly with anti-PD-L1, leading to 40% long-term survivors (Figure 8E). Because CD40a hydrogels are able to mediate these effects with fewer IRAEs than local bolus, hydrogel delivery may be valuable for advancing CD40a-based combination immunotherapy in the clinic.

## 3. Discussion

The most successful passive and injectable carrier for unmodified CD40a to date has been a bolus administration using the viscous montanide oil-in-water adjuvant, which appears to provide moderately sustained delivery of CD40a.^[21]^ Prior efforts to use biomaterials to extend release of CD40a evaluated dextran microparticles, but the material platform drove chronic inflammation that abrogated efficacy.^[22]^ Several modified CD40a approaches have shown promise but introduce added complexity due to the alteration of the antibody. For example, conjugation of CD40a and other agonistic antibodies to liposomal nanoparticles was able to retain drugs at the site of injection and efficiently drain to lymph nodes to bring about superior efficacy and safety,^[23]^ which is consistent with our own findings using a hydrogel carrier. Likewise, a protein-engineered CD40a that binds to extracellular matrix showed promising safety and efficacy.^[24]^ Based on our data, it is likely that the benefits so far noted for local bolus administration can be significantly enhanced by use of injectable hydrogel delivery vehicles. Importantly, unlike many of these previous biomaterials approaches, the PNP hydrogel does not elicit an inflammatory response and does not require chemical modification of the cargo to achieve these benefits.

Using PET imaging, we quantitatively assessed how PNP hydrogel administration improved the pharmacokinetic profile of CD40a compared to local bolus administration. Our data indicated that hydrogels provide a major benefit in terms of increasing drug exposure in target tissues (TME, TdLNs) and reducing off-target exposure, especially in antigen-sink organs like the spleen and distant LNs. Our quantitative analysis of CD40a biodistribution throughout all major tissues provides a comprehensive assessment of the impact of injectable hydrogel delivery on the pharmacokinetics of an immunostimulatory antibody and how that delivery compares to local bolus approaches, which are the current standard for locoregional therapy.

Consistent with the improved pharmacokinetics and biodistribution elicited by hydrogel-based locoregional delivery, we observed administration-dependent changes in the pharmacodynamics of CD40a. In particular, hydrogel administration appeared to attenuate signs of gross toxicity (*e.g.*, weight loss and liver necrosis) compared to local bolus administration. Although these data are in murine models, the pathology for CD40 agonist mediated hepatotoxicity is similar to clinical presentation of this IRAE.^[8a]^ Notably, in our dose reduction study, we identified that hydrogels can safely deliver two times the systemic MTD of CD40a. In contrast, local bolus administration above the systemic MTD continued to present problematic side effects including weight loss and more extensive liver necrosis. Intriguingly, dose-reduction studies with the hydrogel delivery vehicle indicated consistent anti-cancer efficacy of CD40a monotherapy at the ranges tested (100 to 10 μg), indicating an opportunity to pursue dose-sparing strategies. In the context of monotherapy, the efficacy of the lowest dose evaluated (10 μg) appeared to be dependent on hydrogel delivery, which may indicate therapeutic value in the sustained local delivery for CD40a. It is worth noting that administration site and drug delivery vehicles have previously provided dose-sparing effects for infectious disease vaccines.^[25]^ CD40 agonism essentially drives *in situ* vaccination against endogenous tumor antigen,^[26]^ and thus CD40a therapy may exhibit similar dose-sparing effects depending on drug delivery strategy.

One common driver of toxicity for immunostimulatory therapies, including CD40a therapy, is cytokine release syndrome, which is an acute systemic inflammatory condition caused by elevated serum cytokine levels and that can lead to fever and organ disfunction.^[27]^ Analysis of serum cytokine levels revealed that administration method greatly shapes where cytokine induction occurs following CD40a therapy. Unlike local bolus administration, hydrogels can deliver CD40a at a 2-fold higher dose than the systemic MTD without driving higher serum cytokine levels. Notably, at the 2-fold systemic MTD dose, hydrogel delivery led to significantly lower serum levels of key proinflammatory cytokines, including IFNγ, TNFα, IL12, and CXCL10. In particular, the reduction in systemic IL12 may be critical for attenuating CD40-agonist mediated hepatotoxicity.^[28]^ By assessing cytokine induction at varying CD40a dose with both hydrogel and local bolus administration, we identified that drug delivery strategy determined dose-response behavior for peripheral cytokines. In particular, hydrogels reduced systemic cytokine levels by either altering the slope or the elevation of individual cytokine dose-response curves.

While CD40a hydrogels appeared to suppress systemic cytokine induction, these hydrogels mediated significantly higher levels of effector cytokines in the TdLN compared to systemic administration methods. Compared to dose-matched local bolus controls, hydrogel delivery of CD40a induced significantly more IFNγ and CXCL10 in the TdLN. Compared to systemic dosing, local bolus administration also improved the induction of effector cytokines in the TdLN. These results reinforce prior reports that locoregional bolus administration of immunotherapy improves immunogenicity in TdLNs relative to systemic dosing,^[29]^ but our data also indicate that further benefits may be achieved using a biomaterials approach.

We compared the anticancer efficacy of hydrogel and local bolus delivery of CD40a directly in the B16F10 model of melanoma, in both the context of monotherapy and combination therapy. The B16F10 tumors exhibit an immune desert phenotype that is difficult to treat with immunotherapy, and they are known to be resistant to frontline therapies such as immune checkpoint blockade.^[15]^ Notably, CD40a monotherapy has largely been discounted as a viable clinical strategy, where tolerable dosage of the single-agent have yielded disappointing results.^[18]^

Interestingly, a single dose of either 100 or 10 μg CD40a administered *via* hydrogel slowed tumor growth and extended survival, and even presented a low incidence rate (∼10%) of long-term survival. Moreover, we observed an abscopal effect on distant tumors, which indicates relevance for treating metastatic disease. In contrast, no therapeutic benefits were observed with the dose-matched local bolus control at the 10 μg dose. Combining locoregional CD40a therapy with systemic PD-L1 treatment led to synergistic anticancer efficacy, with both hydrogel and local bolus approaches yielding 40% incidence of long-term survivors in the B16F10 model. The similar results between the two delivery approaches may indicate that in combination settings, the 10 μg dose is sufficient for creating an environment receptive to PD-1/L1 blockade. These results indicate that as a monotherapy, potentially more frequent doses of CD40a are needed for curative anticancer efficacy, but in the combination setting, the 10 μg dose used here is sufficient for creating a tumor environment that is receptive to PD-1/L1 blockade.

These results provide a compelling basis for expanding biomaterials-based approaches towards immunomodulation, particularly in comparison to local bolus administration. Prior work has carefully documented the benefits of various strategies for locoregional immunotherapy,^[29–30]^ including for CD40a agonists.^[11c, 21, 26b, 31]^ The most common and well characterized approach has been intratumoral or peritumoral local bolus administration, which has been shown to provide greater immunomodulation in target tissues and reduced toxicity for certain drugs.^[29, 31]^ These benefits are likely attributable to the changes in drug pharmacokinetics due to the brief depot effect that local bolus administration provides, which has been well documented by others,^[23a, 29, 31]^ and is supported by our own data reported here. The ease of local bolus administration is attractive, particularly for more tolerable immunotherapies (*e.g.*, PD-1/L1 blockade), but the depot effect is quite limited and yields underwhelming results for potent immunostimulants such as cytokines and co-stimulatory agonists.^[23a, 32]^ Nevertheless, prior work on local bolus administration of CD40a reported benefits in safety and efficacy,^[21, 31]^ consistent with our own findings. Unfortunately, intratumoral injections of CD40a have recently been studied in the clinic and did not appear to meaningfully alter the therapeutic index of the drug compared to intravenous routes,^[33]^ which may indicate more advanced locoregional strategies, such as the use of injectable hydrogel depot technologies, may be needed to reap the benefits of CD40 agonism. Emerging strategies for multifunctional injectable biomaterials also point towards the possibility of further enhancing the benefits of these systems through the strategic incorporation of drug-binding motifs to further slow release,^[34]^ which could further improve safety and efficacy of immuno-modulatory biomaterials.

Another compelling route for locoregional immunomodulation has been the strategic use of protein engineering to develop immunostimulatory drugs that are either retained or activated in the tumor microenvironment.^[35]^ These sophisticated strategies have clearly demonstrated the power of local immunostimulation and can achieve impressive therapeutic results with minimal toxicity. Yet, the technical difficulty of developing these novel constructs renders this approach inaccessible for many research groups, and often requires the alteration of a clinical drug candidate in ways which may complicate or restart the regulatory process. So, while promising, protein engineering approaches can create a barrier for exploratory preclinical research on the impact of locoregional therapy of different drugs. The injectable hydrogel approach discussed here is drug-agnostic and does not require modification of the encapsulated cargo. Moreover, the synthesis of these hydrogels enables incorporation of drugs through simple mixing under mild conditions.^[36]^ As a result, this approach provides a readily adoptable method to rapidly evaluate diverse locoregional immunotherapy strategies.

## 4. Conclusion

Immunostimulatory drugs such as CD40 agonists, cytokines, TLR agonists, and bispecific antibodies provide a means for repolarizing the immunosuppressive tumor microenvironment toward an immunogenic state. Preclinical and clinical data indicate that incorporating these drugs into combination immunotherapy regimens can significantly improve outcomes for a variety of tumors that fail to respond to either traditional or single-agent immunotherapies.^[6a, 10a, 37]^ Yet, clinical translation of immunostimulatory drugs has been greatly hindered by their propensity toward triggering dose-limiting IRAEs.^[38]^ CD40 agonists, in particular, have had difficulty reaching efficacious doses using conventional systemic administration,^[39]^ and even at these low doses, combination therapies employing these antibodies have observed high rates (47%) of serious adverse effects in pancreatic cancer patients.^[10a]^ Here, we explored a technological solution to this challenge using an injectable supramolecular hydrogel to locally administer CD40 agonist antibodies to the TME and TdLNs, and report that this approach improved the safety profile and anticancer efficacy of this key immunostimulatory drug in both monotherapy and combination therapy settings. Our key findings include the observation that injectable hydrogels provide superior pharmacokinetic benefits over standard local bolus administration. These manifest in the form of increased drug exposure at the injection site and draining lymph nodes and reduced exposure elsewhere in the body. Consistent with these observations, we report that hydrogels substantively impact the off-target toxicity associated with CD40 agonism. Namely, we observed mitigation of acute weight loss and hepatotoxicity despite delivering higher doses of drug. Critically, we observed that hydrogels strongly influence the induction of peripheral and local cytokines, reducing systemic cytokine levels while maintaining elevated cytokine levels in the tumor-draining lymph node. Overall, these data indicate that injectable hydrogels may be valuable tools for implementing CD40a, and potentially other immunostimulants, in both preclinical and clinical contexts.

## 5. Experimental Section/Methods

### Poly(ethylene)_5k_-block-poly(lactic acid)_20k_ synthesis

PEG-b-PLA block copolymers were prepared via organocatalytic ring opening polymerization as follows. DL-lactide (Alfa Aesar) was dissolved in ethyl acetate in the presence of sodium sulfate, and then recrystallized twice. Recrystallized lactide (10 g) was then dissolved in distilled DCM (50 mL; Fisher Scientific) under mild heating and kept under nitrogen. Methoxy-PEG (2.5 g; 5kDa; Sigma Aldrich) was dried under vacuum and heat (90C). Dry methoxy-PEG and distilled 1,8-diazabicyloundec-7-ene (DBU; 75 μL; Sigma Aldrich) were dissolved in distilled DCM (5 mL) and then rapidly added to the lactide solution while under nitrogen and stirred for 8 minutes. The reaction was quenched by addition of acetic acid diluted in acetone (1-2 drops in 0.5 mL). The product was precipitated three times in 50:50 hexane/ether solution, after which it was dried under vacuum. Size and uniformity of the product was confirmed using GPC. Briefly, GPC samples were prepared by dissolving PEG-b-PLA (5 mg) in DMF (1 mL) and filtering through 0.2-micron nylon syringe filter (Tisch Scientific). M_n_ and M_w_ were determined after passing through two size exclusion chromatography columns (Resolve Mixed Bed Low DVB, ID 7.8 mm, Mw range 200–600,000 g mol^−1^ (Jordi Labs)) in a mobile phase of N,N-dimethylformamide (DMF; Sigma Aldrich) with 0.1 M LiBr at 35 °C and a flow rate of 1.0 mL min^−1^ (Dionex Ultimate 3000 pump, degasser, and autosampler (Thermo Fisher Scientific)) using PEG standards (American Polymer Standards Corporation).

### PEG-b-PLA nanoparticle synthesis

PEG-b-PLA (50 mg) was dissolved in acetonitrile (1 mL) and added dropwise to 10 mL of rapidly stirring Millipore water to form 30 nm nanoparticles via nanoprecipitation. Size and uniformity of the nanoparticles were confirmed using a Wyatt DynaPro dynamic light scattering instrument. The dilute nanoparticle product (11 mL) was concentrated using a 10 kDa Amicon spin filter (4500 rcf for 1 hour). The concentrated nanoparticle was diluted to a 20% w/w solution using 1x PBS, then stored at 4°C until used.

### Dodecyl-modified hydroxypropylmethylcellulose (HPMC-C_12_) synthesis

Hypromellose (1 g; Sigma Aldrich) was dissolved in anhydrous N-Methylpyrrolidone (NMP; 40 mL; Sigma Aldrich) and heated to 80°C in a PEG bath. Dodecyl isocyanate (125 μL; Sigma Aldrich) was diluted in anhydrous NMP (5 mL) and added dropwise to the Hypromellose solution while rapidly stirring. N,N-Diisopropylethylamine (HUNIGS; 10 drops; Sigma Aldrich) was added dropwise to the reaction solution while rapidly stirring. The heat bath was turned off, and the reaction was allowed to continue overnight while stirring. The polymer was precipitated in acetone (600 mL) and then dissolved in Millipore water (∼40 mL) prior to being dialyzed (3.5 kDa MWCO; Spectrum Labs) for 4 days. Pure HPMC-C_12_ was then lyophilized and dissolved in 1x PBS (Thermo Fisher) to produce a 6% w/v solution, which was stored at 4°C until used.

### Polymer-nanoparticle hydrogel synthesis

PEG-PLA NP, HPMC-C_12_, and antibody solutions were mixed using a dual syringe technique as described previously.^[36]^ Gels were composed of 50% v/v of PEG-PLA NP solution (prepared to 20% w/w as described above), 33.4% v/v of HPMC-C_12_ (prepared to 6% w/v as described above), and 16.6% v/v of antibody solution (using the desired concentration for a given dosage). PEG-PLA NPs were mixed with the antibody solution and loaded into an appropriately sized syringe. HPMC-C_12_ was loaded into a separate syringe and connected to an autoclaved elbow connector. HPMC-C_12_ solution was gently pushed into the elbow connector, and then the NP/antibody syringe was connected carefully to minimize air bubbles within the dual-syringe system. Once securely connected, the solutions were vigorously mixed by alternating depression of the connected syringes (100 cycles minimum mixing) to form the PNP hydrogel. After mixing, one syringe was depressed fully to transfer the gel to the other syringe, which could then be detached and stored at 4°C until used.

### Rheological characterization of hydrogels

Mechanical properties were assessed using a TA Instruments Discovery HR-2 torque-controlled rheometer fitted with a Peltier stage. Approximately 0.5 mL of hydrogel was deposited onto a serrated 20 mm plate geometry for all measurements, and all measurements were carried out at room temperature. Initial gap size was set to 0.5 microns and adjusted to adequately load the geometry. Dynamic oscillatory frequency sweep measurements were performed with a constant torque ranging from 0.1 to 100 s^−1^. Steady shear flow rheology was performed from 0.1 to 10 s^−1^. Step-shear experiments were performed by alternating between high (10 s^−1^) and low (0.1 s^−1^) shear rates for at least two full cycles.

### Mammalian cell culture

Murine B16F10 melanoma was purchased from ATCC. MC38 colorectal cancer cells were a gift provided by the Davis Lab at Stanford. B16F10 cells were maintained using DMEM media (Thermo Fisher) supplemented with 10% fetal bovine serum (Novus Biologicals) and 1% penicillin-streptomycin (Thermo Fisher) prior to tumor inoculation. MC38 cells were cultured using DMEM media supplemented with 10% fetal bovine serum, 1% penicillin-streptomycin, and 10 mM 4-(2-hydroxyethyl)-1-piperazineethanesulfonic acid (HEPES; Sigma Aldrich). All cell lines were tested for mycoplasma using the MycoAlert Microplasma Kit (Lonza) prior to tumor inoculation.

### Murine tumor models

All animal studies were performed following Stanford’s IACUC guidelines and protocols with the approval of the Stanford Administrative Panel on Laboratory Animal Care (APLAC-32947). MC38 flank xenografts were generated on 7-week old female C57BL/6 mice by injecting a 100 μL bolus of MC38 cells (5×10^6 cells/mL in PBS) subcutaneously above the right hind leg. B16F10 melanoma models were generated similarly; 7-week old female C57BL/6 mice were injected with a 100 μL bolus of B16F10 cells (3×10^6 cells/mL in PBS) subcutaneously above the right hind leg. For the metastatic model of B16F10, tumors were inoculated as described, but on both right and left flanks. Cells were maintained and expanded prior to inoculation as described above. Following tumor inoculation, mice were monitored for palpable tumors. Once tumors grew to 2.5 mm or larger, mice were randomized into a treatment cohort and dosed within 24 hours. For models of established tumors, mice were not treated until tumors grew larger than 3.5 mm. Mouse weight was tracked to assess acute toxicity, with euthanasia criteria for sudden losses of more than 20% of body mass relative to the start of treatment. Tumor burden was tracked using caliper (Mitutoyo) measurements to calculate total tumor area (L x W); mice were euthanized once total tumor burden grew to 150 mm^2^ or larger.

### Peritumoral injection of treatments

Local therapies were administered using a peritumoral injection technique. This technique consisted of tenting the skin near the tumor to allow for a syringe to be inserted into the subcutaneous space. The needle of the syringe was guided through the subcutaneous space until the needle was within a few millimeters of the tumor, this must be done carefully to avoid piercing the skin which would allow leakage of saline bolus injections. Once the needle is in position, the treatment was injected into the space around the tumor. Following injection, the needle was left in place for roughly 10-15 seconds. This is to allow for any compressed hydrogel to relax and finish extruding from the needle. This also allows the local bolus to begin to spread out and minimizes leakage of the dose upon removal of the needle.

### Positron emission tomography pharmacokinetics study

CD40a (clone FGK4.5; Bio X Cell) were radiolabeling as described previously.^[40]^ Briefly, CD40a were conjugated to the bifunctional chelator p-SCN-Bn-Deferoxamin via NHS addition chemistry. Antibodies were buffer exchanged to pH 9 PBS using Vivaspin 2 desalting columns. Antibody was mixed with p-SCN-Bn-Deferoxamin (Macrocyclics B-705) at a 1:10 molar ratio, and incubated for 1 hour at 37°C while stirring (450 RPM) on a Thermomixer C. CD40a-DFO was buffer exchanged to pH 7.4 PBS after incubation. All bioconjugation was performed using low binding plasticware, when possible. ^89^Zr-oxalate was obtained from 3D Imaging (Little Rock, AR) and upon arrival was diluted to 100 μL using Traceless water and titrated to a pH of 7.1-7.9 using 1 M sodium carbonate. 1 mCi of ^89^Zr-oxalate was added to 300 μg CD40a-DFO. The mixture was incubated for 1 hour at 37C, and labeling was confirmed using iTLC. The radiolabeled antibody was then purified using a desalting column. PNP hydrogels were prepared, as described above, with 80% cold antibody and 20% hot antibody. B16-tumor bearing mice were injected with a total of 150 μL of gel containing 100 μg total of CD40a (20 μg radiolabeled). Mice were imaged using an Inveon microPET/CT two hours after injection, and then every 24 hours out to 12 days post-injection. Data was exported to the Inveon Research Workspace, where regions of interest for specific tissues were defined based on CT images. ROI data, in nCi/cc, were transformed to %ID/g using the following formula:

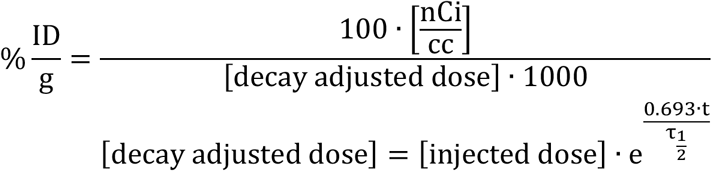

### Luminex cytokine assay and statistical analyses

Mice bearing B16F10 melanoma flank xenografts were treated with CD40 agonist antibodies via peritumoral hydrogel or via dose-matched local bolus injections. Additional controls included saline treated, isotype antibody (BE0089; BioXCell) loaded hydrogel, and 5 ug of CD40a delivered systemically (IP injection). 24 hours after treatment, 90 uL of blood were collected via tail vein bleeds and centrifuged using Sarstedt Z-gel microtubes (ThermoFisher) to isolate serum. Serum was analyzed by Stanford’s Human Immune Monitoring Center (HIMC) using a 40-plex murine Luminex cytokine panel. 72 hours after treatment, mice were euthanized and tumor draining inguinal lymph nodes were excised and processed for analysis. TdLNs were weighed and then 10 mL of lysis buffer (Cell Extraction Buffer FNN0011 with 1x Halt protease inhibitor, ThemoFisher) was added per 1 g of tissue. Tissues were ground using a Dounce Tissue Grinder, after which the sample was collected in protein lo-bind Eppendorf tubes and centrifuged at 10,000 RCF for 5 minutes to pellet tissue debris. Supernatants were collected into separate protein lo-bind Eppendorf tubes and stored at −80°C. Tissues/homogenates were kept on ice throughout processing. Samples were thawed and total protein content was analyzed using a Pierce BCA assay, and sample protein content was normalized when possible to 2 mg/mL, diluting with additional lysis buffer. TdLN homogenates were provided to the HIMC for analysis by a 40-plex murine Luminex cytokine panel.

Analyses of the data were carried out through the statistical consultation service provided by the HIMC. Briefly, data processing began by removing plate and nonspecific-binding artifacts from the MFI data for each protein.^[20]^ In this process, MFI values for each specimen and protein were averaged across replicate wells. Let Y_ij_ represent a mean MFI value from the i^th^ specimen on the j^th^ protein.

For the serum specimens, we regressed Y_ij_ on cohort:

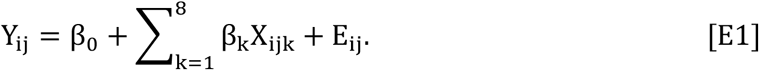

X_ijk_ is 1 if the observation is from the k^th^ treatment cohort and 0 otherwise, with the convention that X_ijk_ = 0 for all k for the untreated cohort, thereby making β_0_ the mean for untreated and each Β_k_, k ∈ {1, …, 8}, the difference for the k^th^ treated cohort’s mean from the untreated mean. The eight alternative hypotheses of interest were β_k_ ≠ 0, k ∈ {1, …, 8}. E_ij_ is residual (i.e., unexplained) error.

For the lymph-node specimens, we regressed Y_ij_ on cohort and total protein content:

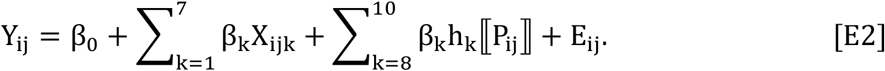

Notation is similar to [E1]. The untreated condition is excluded (due to its absence from one plate); so, β_0_ is the mean of the 5 mg systemic bolus, against which all other treatment cohorts are compared. Also, now is included a piecewise constant basis expansion in two knots for total protein content P_ij_.^[41]^ This spline allowed for a nonlinear relationship between total protein content and MFI and with a minimum of regression parameters because sample size was modest (n = 45). Knots for the spline were placed at the values of 1 and 2 of total protein content.

Regression models [E1] and [E2] were estimated using generalized maximum entropy estimation on a wide prior support of −1,000 to +1,000.^[42]^ Distributions of the Y_ij_ vary among proteins and GMEE does not make any assumptions about the sampled distribution.

False discovery rate (FDR) was controlled at 5% using Benjamani et al. per Kim and van de Wiel.^[43]^ For lymph nodes and serum, the FDR was controlled across all 48 cytokines and 7 means comparisons or 48 ⨉ 7 = 336 hypothesis tests because comparisons were unplanned,^[44]^ given that 5 mg systemic bolus was chosen as the comparator based upon inspection of some of the data. Cluster analyses began by using the data that had been corrected for plate and nonspecific binding artifacts. Let Y_ij_ represent the mean corrected MFI value from the i^th^ specimen on the j^th^ protein. For serum, the Y_ij_ values constitute the data set input to the cluster analyses.

The lymph node data also required correction for total protein. We regressed Y_ij_ on total protein content:

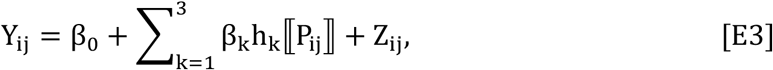

using the same piecewise constant basis expansion in two knots for total protein content P_ij_ as defined in [E2]. We fit [E3] using ordinary least squares.^[45]^ For lymph nodes, the data corrected for total protein, the residual Z_ij_ values, constituted the data set input to the cluster analyses.

For serum and for lymph nodes, hierarchical sparse clustering was performed.^[46]^ Separately for each data set, five hundred random permutations were performed to select the tuning parameter.

### Treatment efficacy studies

To evaluate the ability for locoregional CD40 therapy to induce an abscopal effect, dual-flank B16F10 tumors were inoculated as described above. Mice were randomized into treatment groups once tumor size grew larger than 2.5 mm in diameter, and were treated the following day with peritumoral injection of 100 μL of hydrogel containing either 100 μg of anti-CD40 or anti-PD-1 (clone RMP1-14; Bio X Cell) antibodies. As a negative control, empty hydrogels (also 100 μL) were also injected. To simulate frontline immunotherapy, one treatment group was systemically dosed (via intraperitoneal administration) with 250 μg of anti-PD-1 antibody each week, up to three times, formulated in 200 μL of sterile PBS. Tumor monitoring and euthanasia criteria are as described above.

To evaluate the ability for low dose locoregional CD40 therapy to exert antitumor effects, mice bearing a single established B16F10 tumor were generated as described above. Once tumors were larger than 3.5 mm in diameter, mice were randomized into treatment groups and allocated to cages so that each treatment group was equally represented per cage, which allowed us to use cage covariate data as a blocking variable in our statistical analysis. The following day, mice were treated with either i) peritumoral injection with 100 μL hydrogel containing 10 μg of anti-CD40 agonist antibody, ii) peritumoral injection with 100 μL hydrogel containing 10 μg of isotype control antibody (BE0089; Bio X Cell), or iii) local bolus administration of 10 μg of anti-CD40 agonist antibody in 50 μL sterile PBS. Tumor monitoring and euthanasia criteria are as described above. To evaluate combination therapy conditions, mice were generated as stated above and treatment regimens were supplemented PD-L1 (clone 10F.9G2; Bio X Cell) or isotype (BE0090; Bio X Cell) antibody. More specifically, combination therapy mice were systemically dosed (via intraperitoneal administration) with 250 μg of anti-PD-1 antibody every 2 days, up to three times, formulated in 200 μL of sterile PBS. The PD-L1 antibody therapy began 2 days after locoregional CD40a treatments.

Tumor growth and overall survival were assessed using general linear models with the SAS statistical software. For tumor growth, the Mixed Procedure method was used with input variables consisting of time, specimen identifier, treatment, cage (when appropriate), and the natural log transformed tumor area. For overall survival analysis, the LIFEREG procedure was used with the input variables consisting of treatment, days survived, event (to identify censored data), and cage. Treatment was always used as a blocking factor, and when permitted by the study design, specimen and cage were used as blocking factors. False discovery rate (FDR) was controlled at 5% using the Bejamini and Hochberg method.

## Conflicts of Interest

The authors declare that they have no competing interests.

## Acknowledgements

The authors would like to thank the Stanford Core Facilities that were central to the completion of this work. We extend our gratitude to Dr. Tyson H. Holmes and Dr. Yael Rosenberg-Hasson at the Stanford Human Immune Monitoring Center. We would also like thank Dr. Kelly D. Moynihan for helpful discussion and advice on this work. This research was financially supported by the American Cancer Society (RSG-18-133-01-CDD) and the Goldman Sachs Foundation (administered by the Stanford Cancer Institute, SPO# 162509). S.C. is supported by the National Cancer Institute of the National Institutes of Health under Award Number F32CA247352. E.C.G. would like to thank the NIH Cell and Molecular Biology Training Program (T32 GM007276) E.L.M. would like to thank the NIH Biotechnology Training Program (T32 GM008412). A.K.G. is thankful for a National Science Foundation Graduate Research Fellowship and the Gabilan Fellowship of the Stanford Graduate Fellowship in Science and Engineering. O.M.S. is thankful for a National Science Foundation Graduate Research Fellowship. C.S.L. is supported by the National Defense and Science Engineering Graduate (NDSEG) Fellowship Program.

## Notes

### Competing Interest Statement

S.C., E.C.G., A.K.G., and E.A.A. are on various granted or pending patents reporting the hydrogel technology described in this article (U.S. Patent Application No. 63/094716; International Patent Cooperation Treaty Application No. PCT/US2019/054070).

