## Supplemental Materials for "Injectable Nanoparticle-Based Hydrogels Enable the Safe and Effective Deployment of Immunostimulatory CD40 Agonist Antibodies"

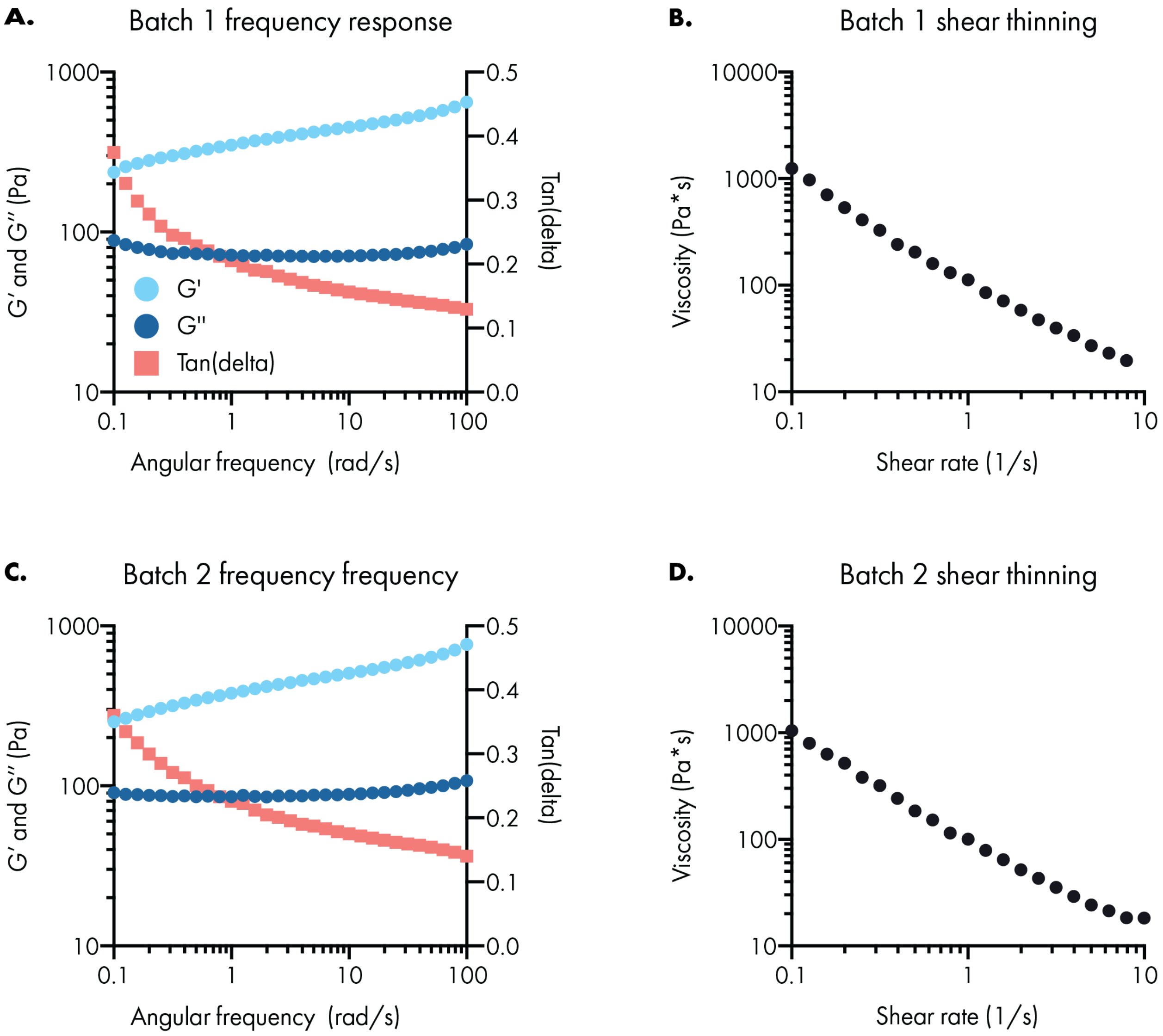


**Figure S1.** Additional rheology data for PNP hydrogels.

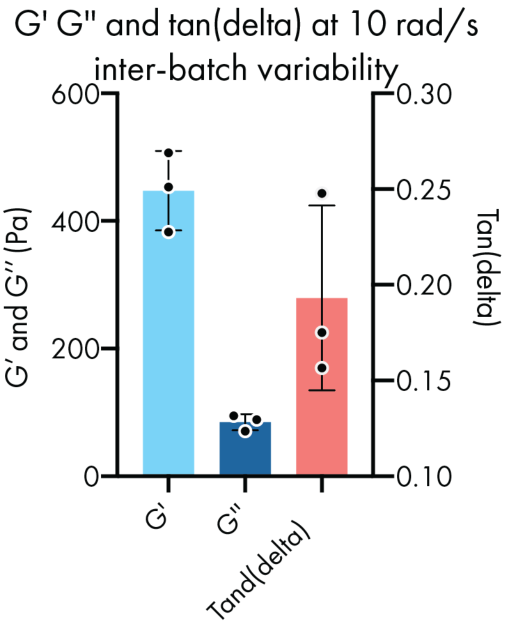


**Figure S2.** Inter-batch variability of the rheological properties for PNP hydrogels.

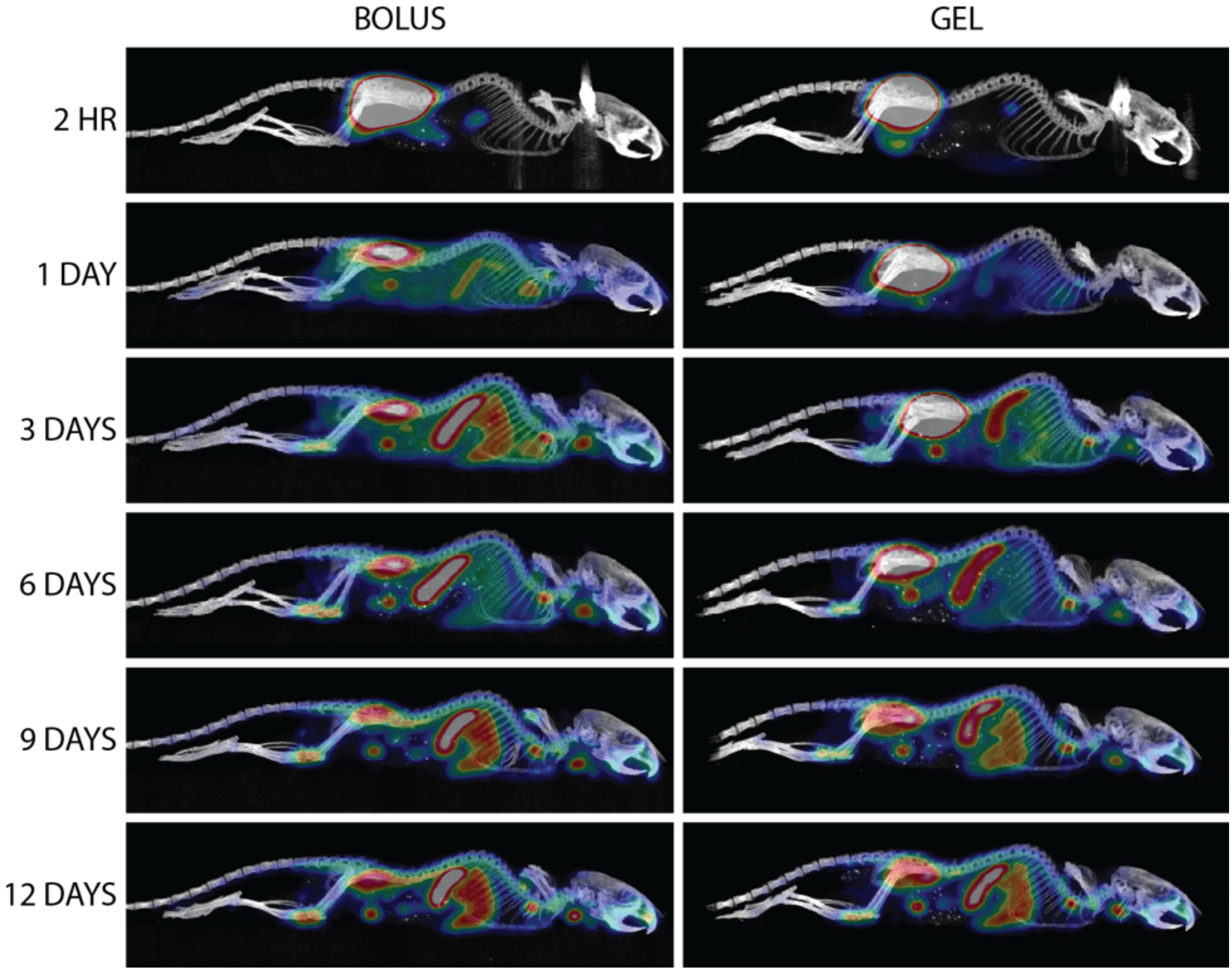


**Figure S3.** Representative PET images comparing CD40a biodistribution over the course of 12 days after administration via local bolus or hydrogel.

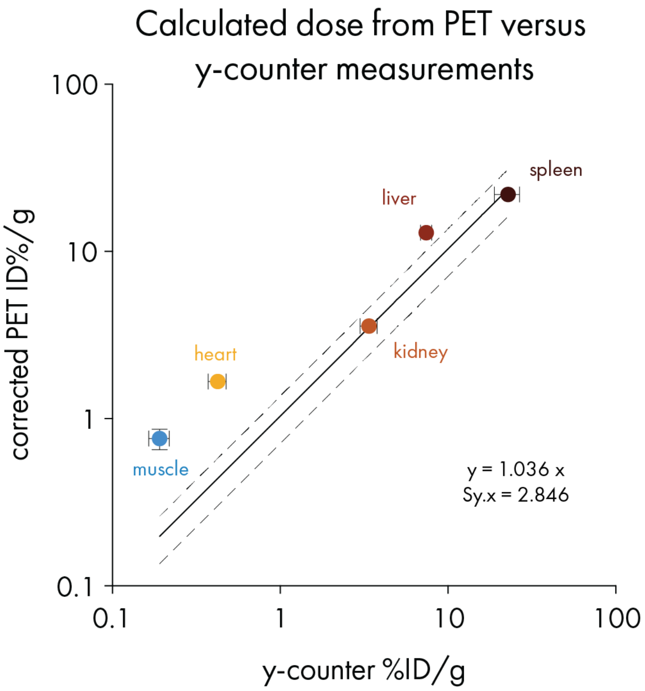


**Figure S4.** Biodistribution data calculated from PET scans compared to values measured from explanted organs using a gamma counter.

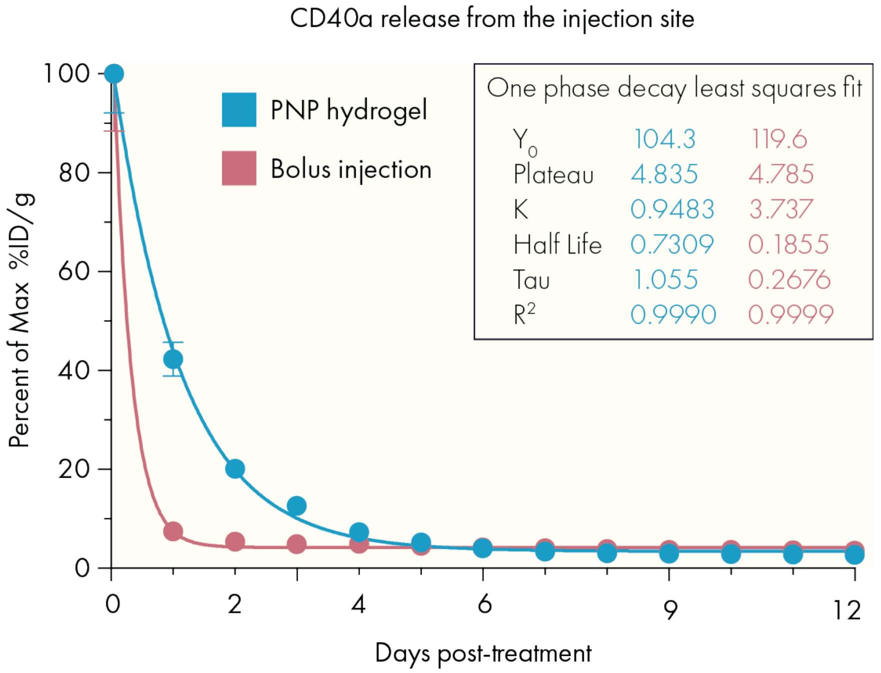


**Figure S5.** Quantified CD40a concentration over time curves at the injection site from PET imaging.

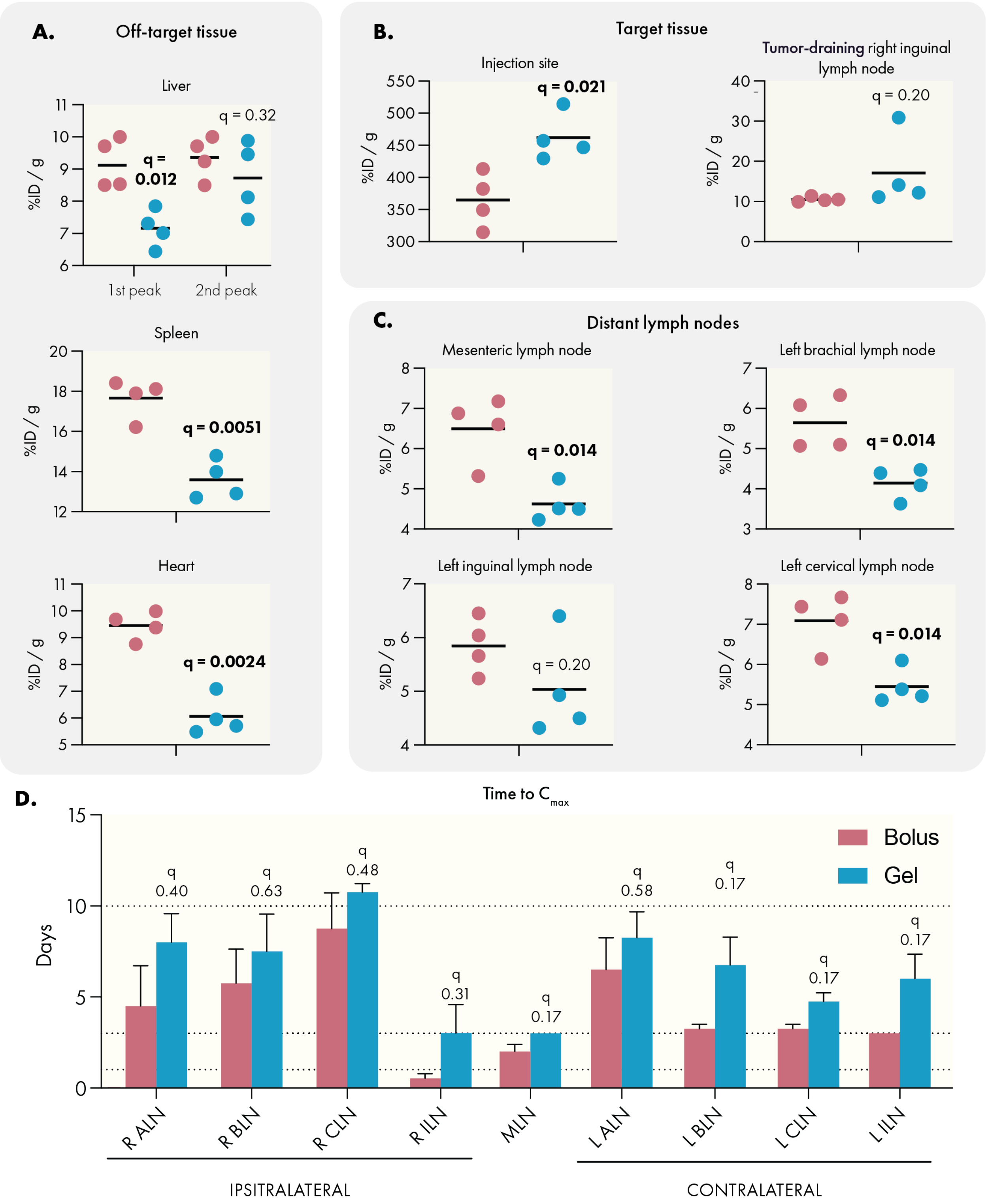


**Figure S6.** Additional pharmacokinetic metrics for CD40a delivery, calculated from PET biodistribution study. (A) $C_{max}$ values for off-target organs. (B) $C_{max}$ values for target tissues. (C) $C_{max}$ values for distant lymph nodes. (D) Mean time to $C_{max}$ for lymph nodes, error bars represent SEM. N = 4 for each treatment.

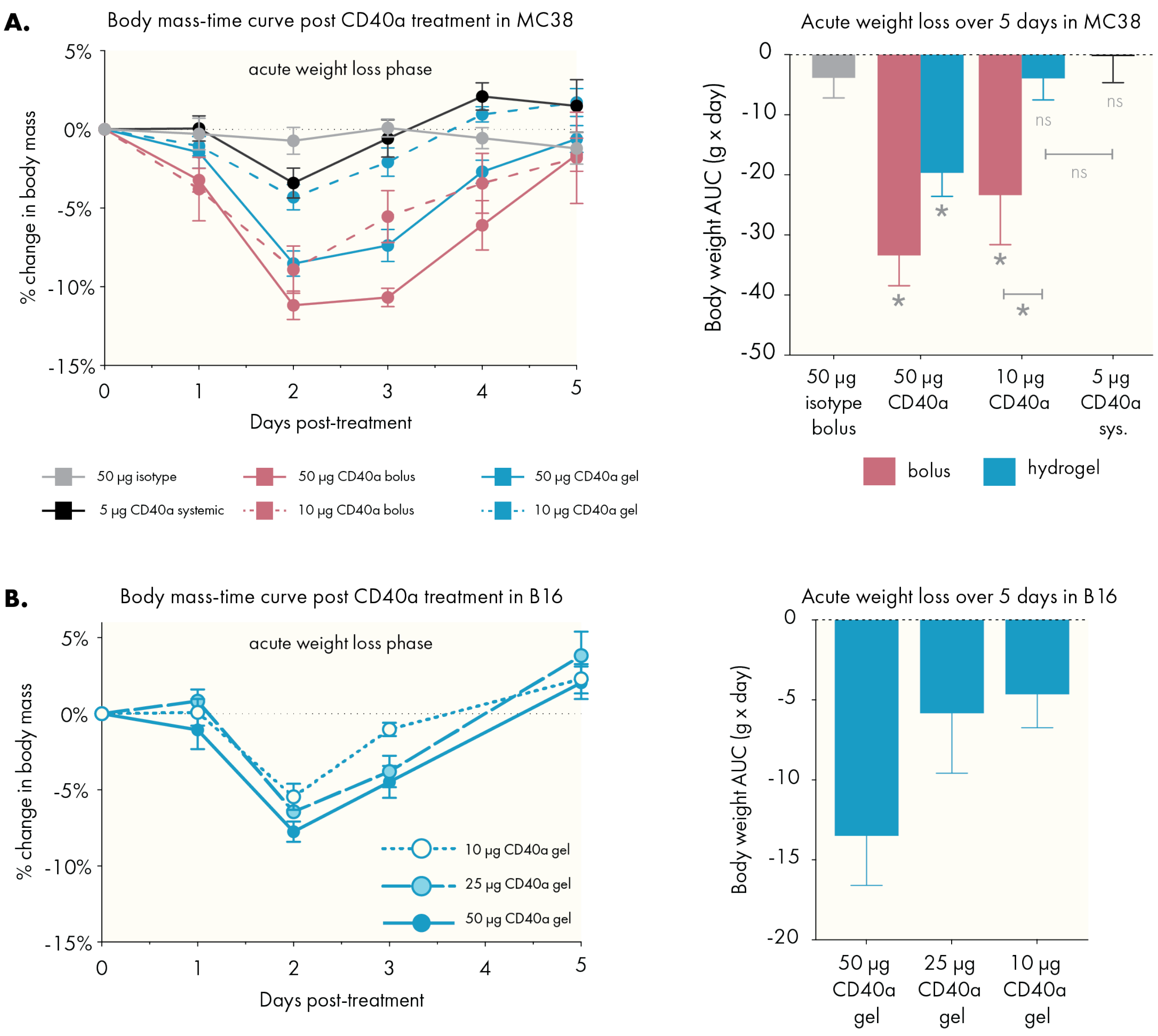


**Figure S7.** Body weight-time curves for MC38 and B16 studies in mice.

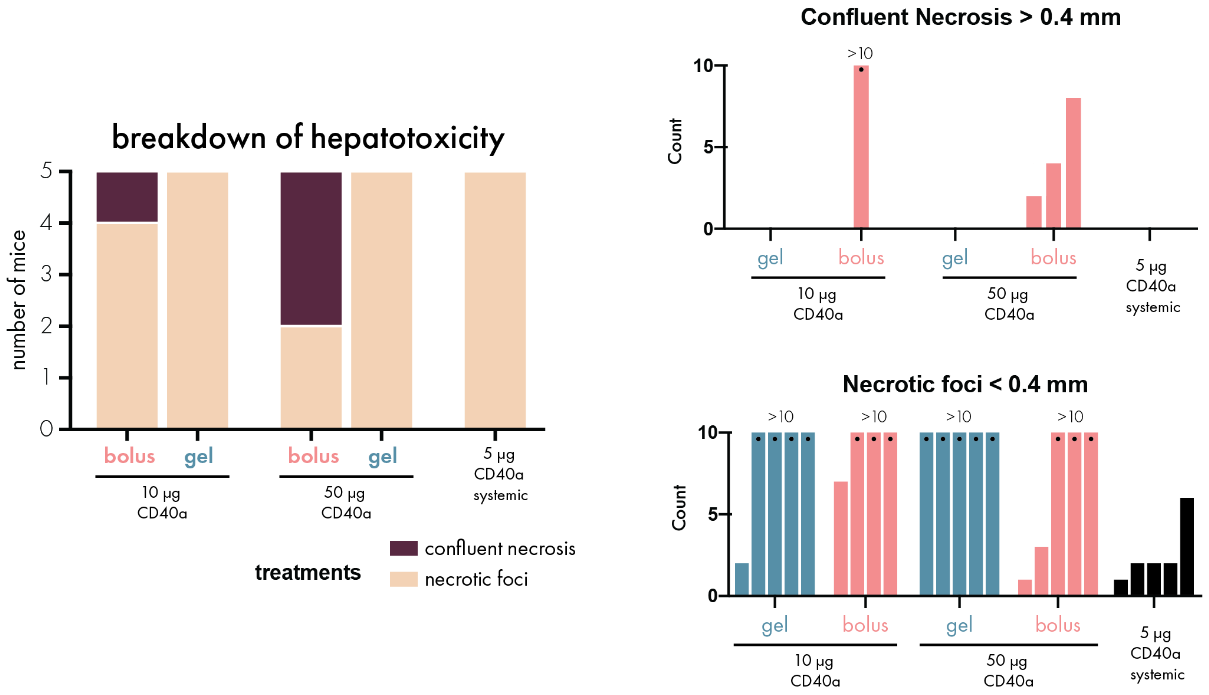


**Figure S8.** Quantitative breakdown of liver toxicity determined from liver histology 72 hours after treatment with CD40a.

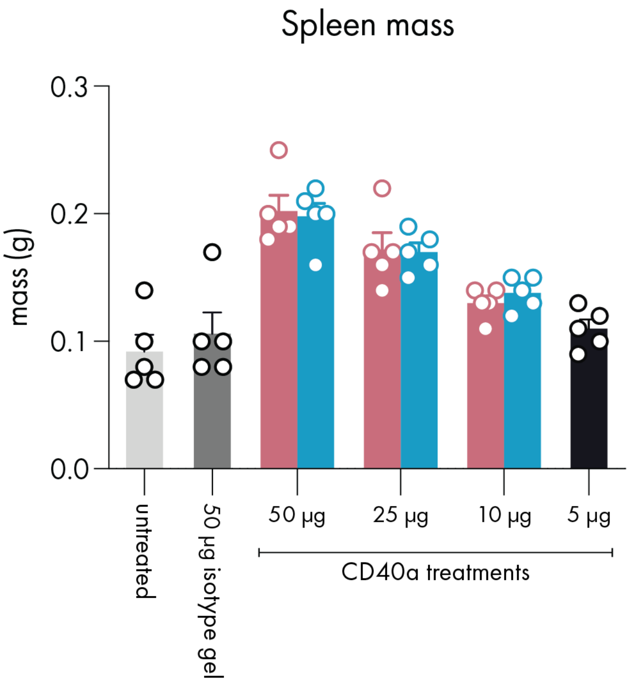


**Figure S9.** Spleen mass at different doses in gel or bolus 72 hours after treatment.

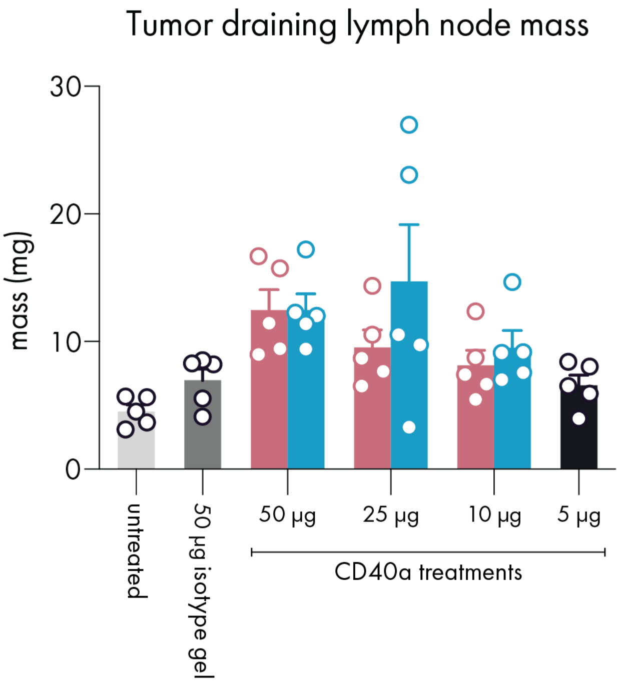


**Figure S10.** Lymph node mass following different doses of CD40a in gel or bolus, 72 hours after treatment.

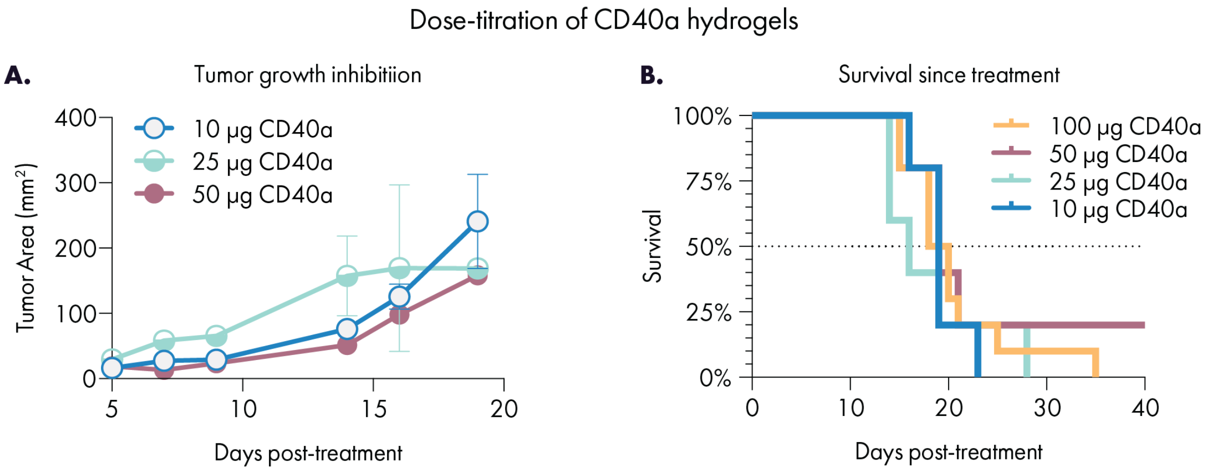


**Figure S11.** Growth and Survival curves for dose titration in B16F10 using hydrogel delivery vehicles. N = 5 for 10, 25, and 50 μg groups and n = 10 for the 100 μg group.

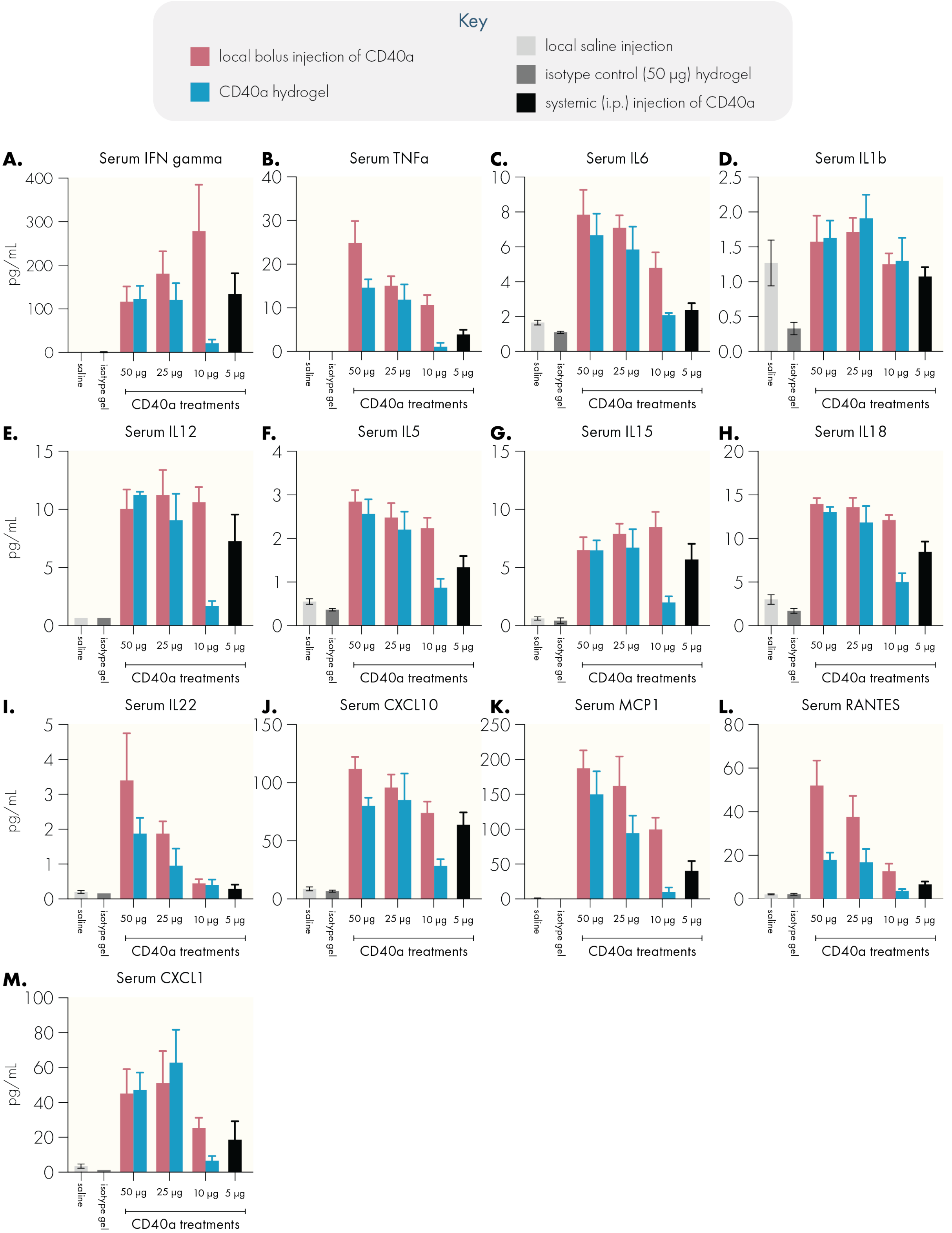


**Figure S12.** Conversion of serum Luminex MFI signals to pg/mL serum concentrations based on standard curves for selected cytokines. For reasons discussed in the main text, we provide these concentrations for comparison to other literature but center our analyses on more accurate assessments based on the corrected fluorescence measurements shown below.

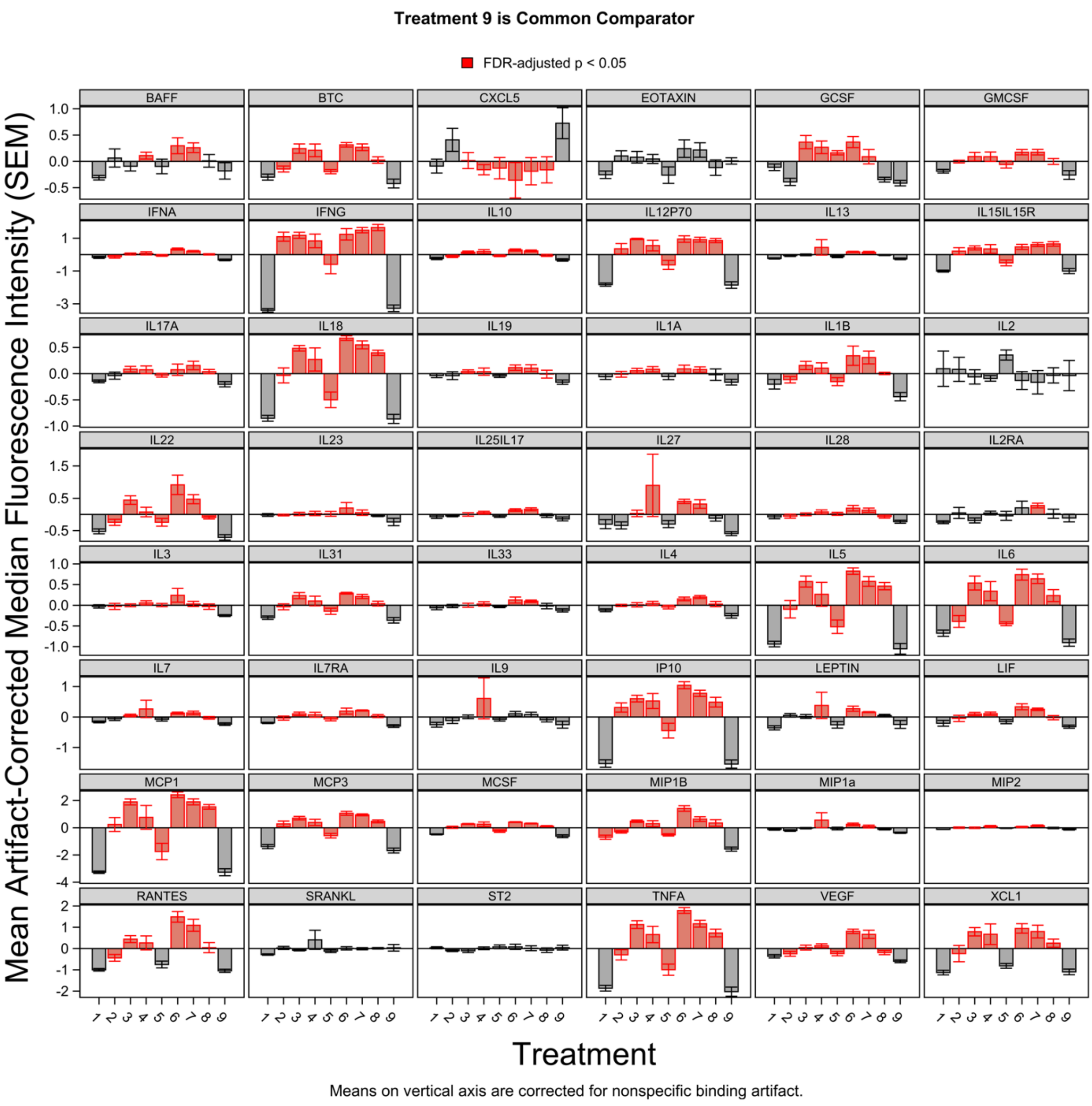


**Figure S13**. Full artifact-corrected Luminex dataset for serum cytokine levels 24 hours after treatment. Treatment key: **1** = 50 µg isotype control; **2** = 5 µg CD40a systemic dose (maximum tolerated systemic dose); **3** = 50 µg CD40a in hydrogel; **4** = 25 µg CD40a in hydrogel; **5** = 10 µg CD40a in hydrogel; **6** = 50 µg CD40a as local bolus; **7** = 25 µg CD40a as local bolus; **8** = 10 µg CD40a as local bolus; **9** = untreated.

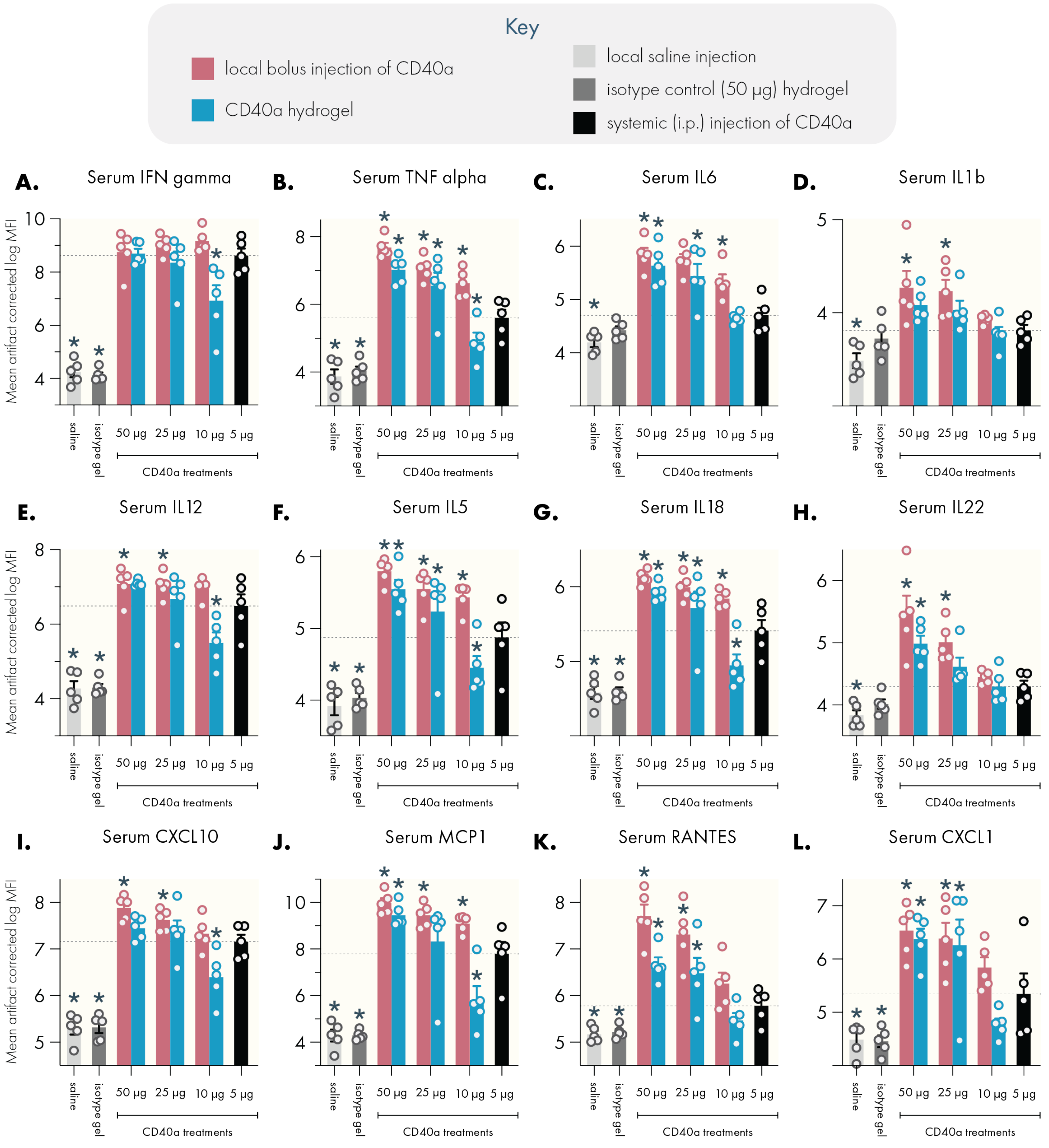


**Figure S14.** Highlighted artifact-corrected serum cytokine data. Dotted line indicates mean value corresponding to the 5 µg systemic dose (maximum tolerated systemic dose). N = 5 for all groups, data represented as mean and SEM. * Denote a significant difference ($p < 0.05$) from the MTD systemic dose. Multiple testing error was controlled using the FDR approach (Q = 5%).

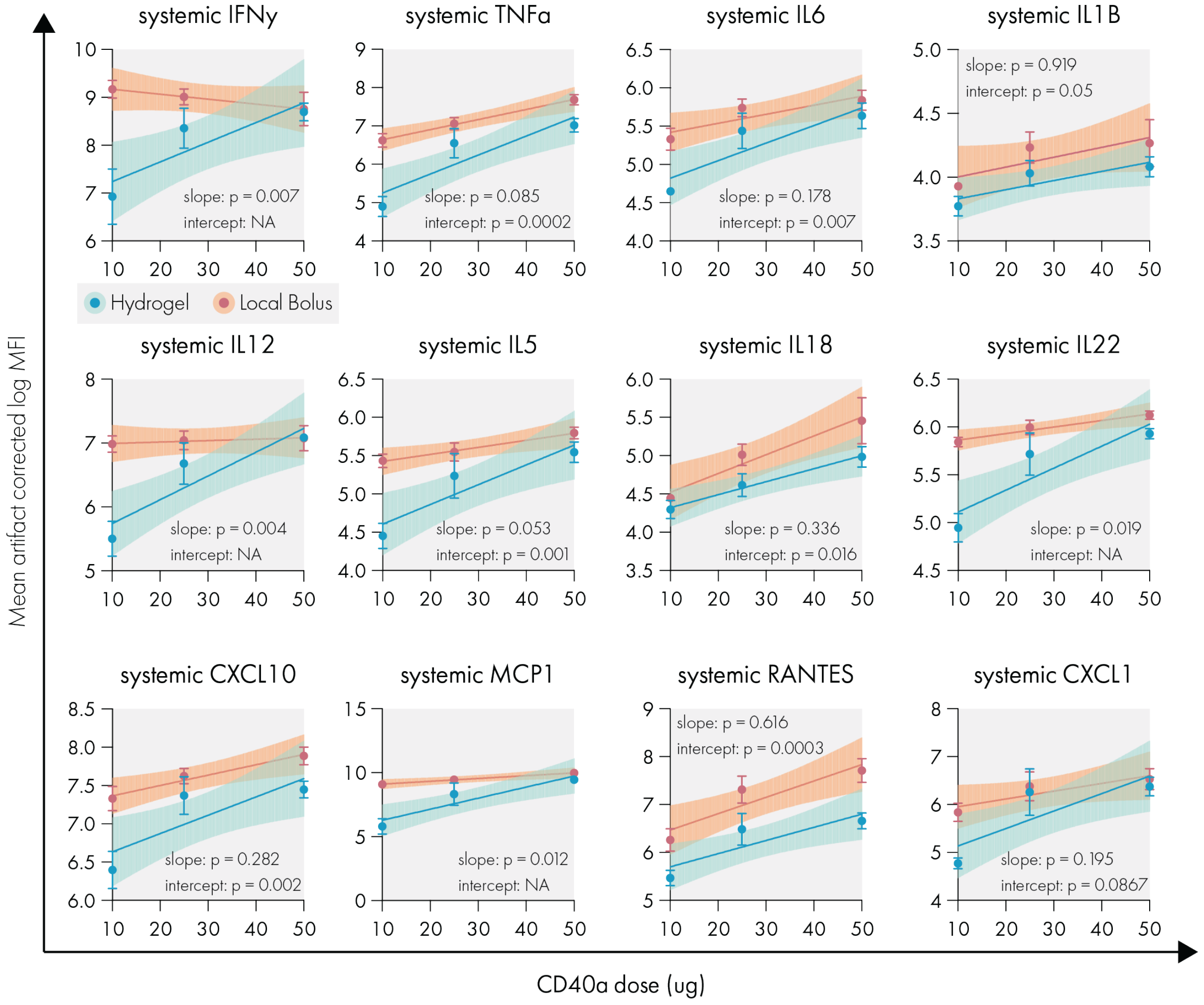


**Figure S15.** Simple linear regression, with no constraints, of dose-serum cytokine response curves for individual cytokines. Shaded area represents 95% confidence interval, error bars indicate SEM. Statistical comparisons performed using built-in analysis in the linear regression functionality of GraphPad Prism.

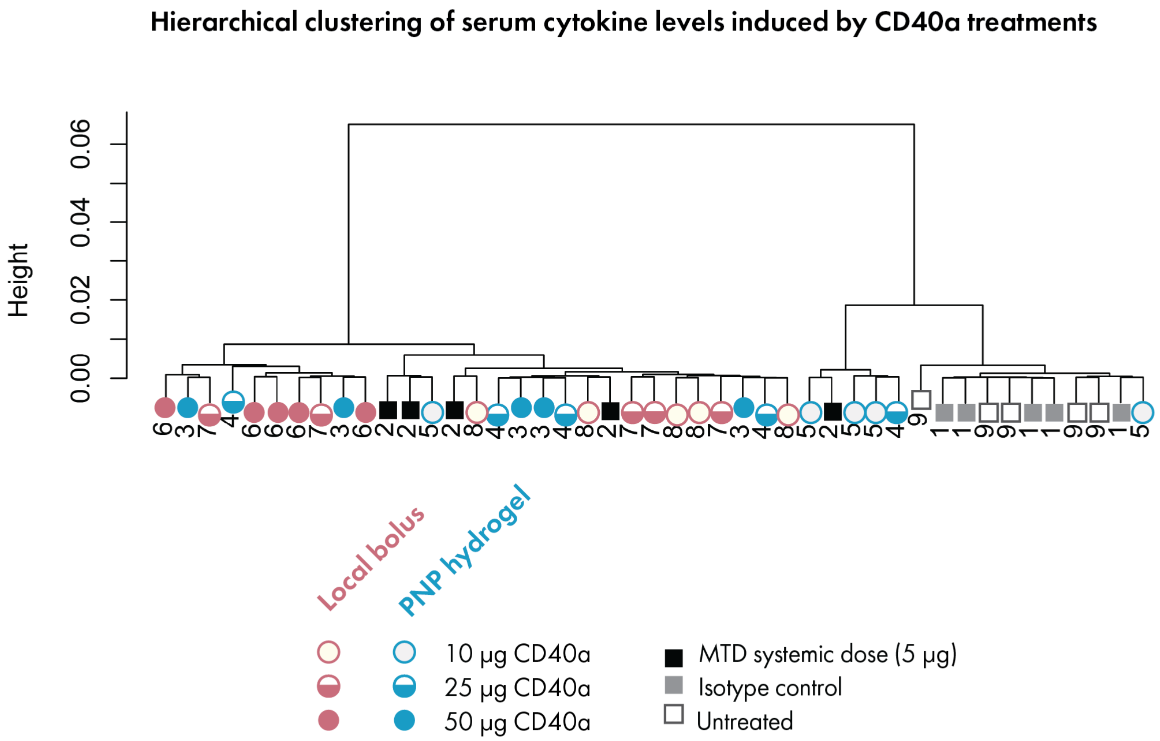


**Figure S16.** Serum cytokine response dendrogram.

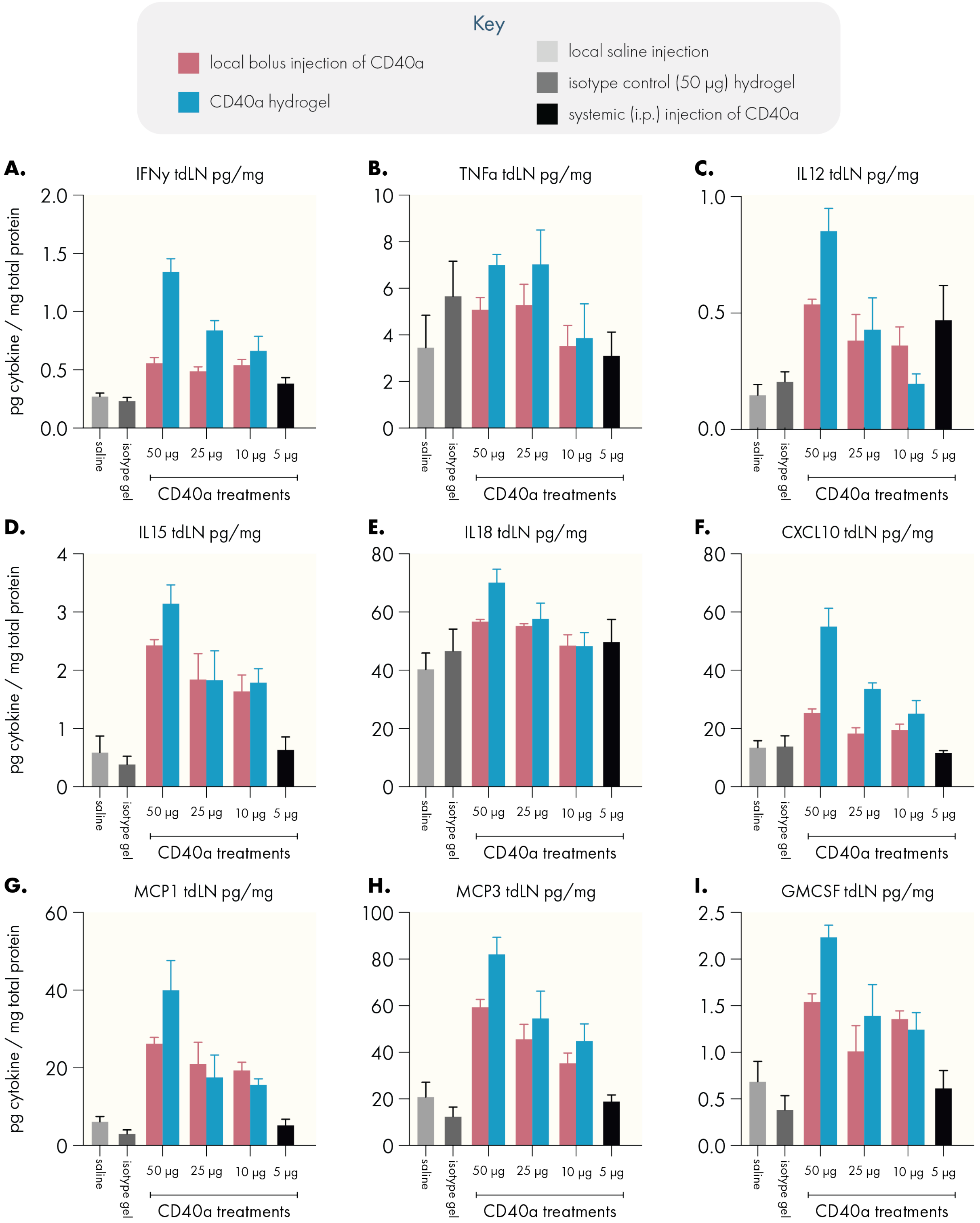


**Figure S17.** Conversion of serum Luminex MFI signals to pg/mg total protein based on standard curves for selected cytokines. For reasons discussed in the main text, we provide these concentrations for comparison to other literature but center our analyses on more accurate assessments based on the corrected fluorescence measurements shown below.

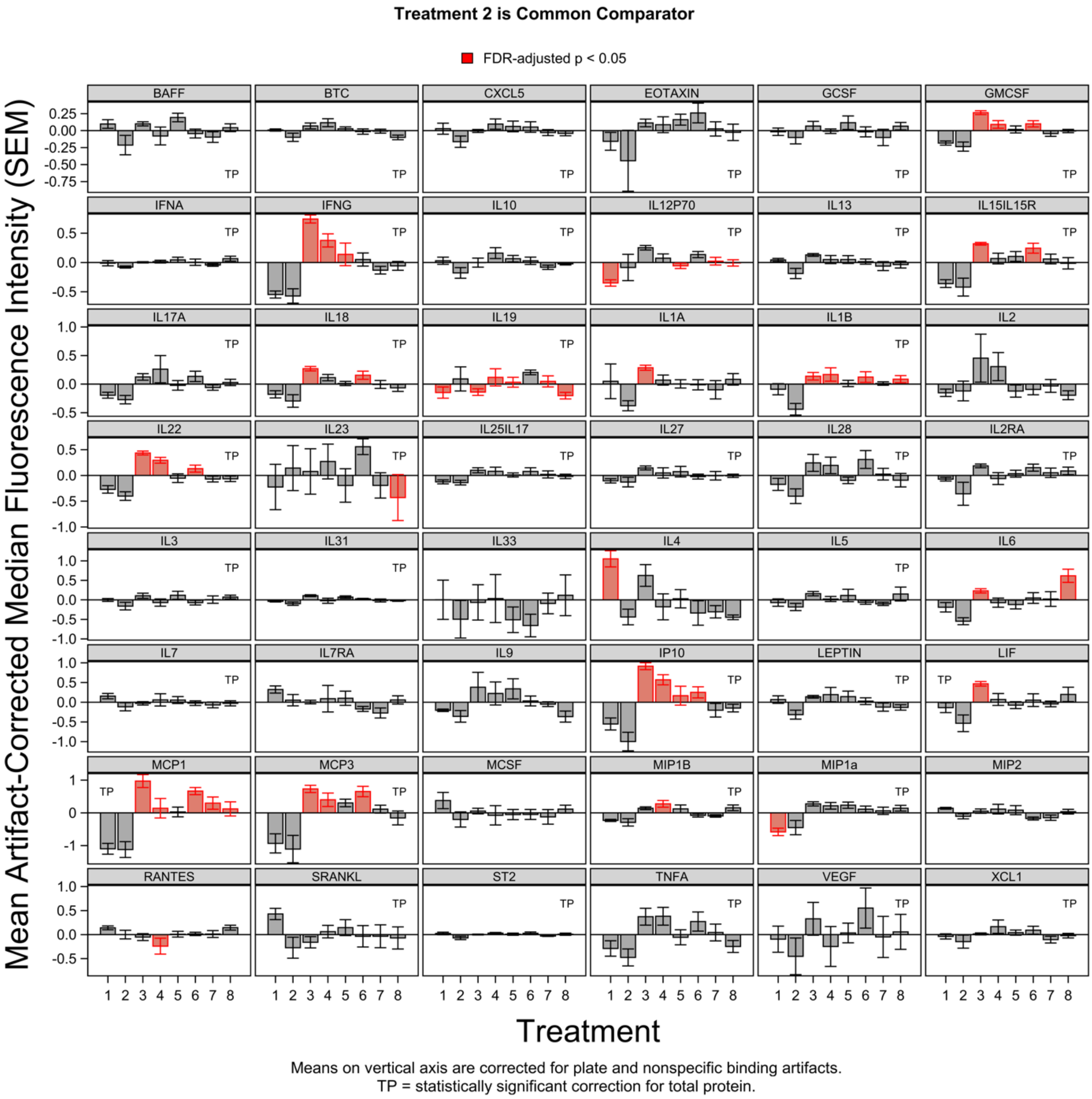


**Figure S18.** Full artifact-corrected Luminex dataset for tumor-draining lymph node cytokine levels 3 days after treatment. Treatment key: **1** = 50 µg isotype control; **2** = 5 µg CD40a systemic dose (maximum tolerated systemic dose); **3** = 50 µg CD40a in hydrogel; **4** = 25 µg CD40a in hydrogel; **5** = 10 µg CD40a in hydrogel; **6** = 50 µg CD40a as local bolus; **7** = 25 µg CD40a as local bolus; **8** = 10 µg CD40a as local bolus.

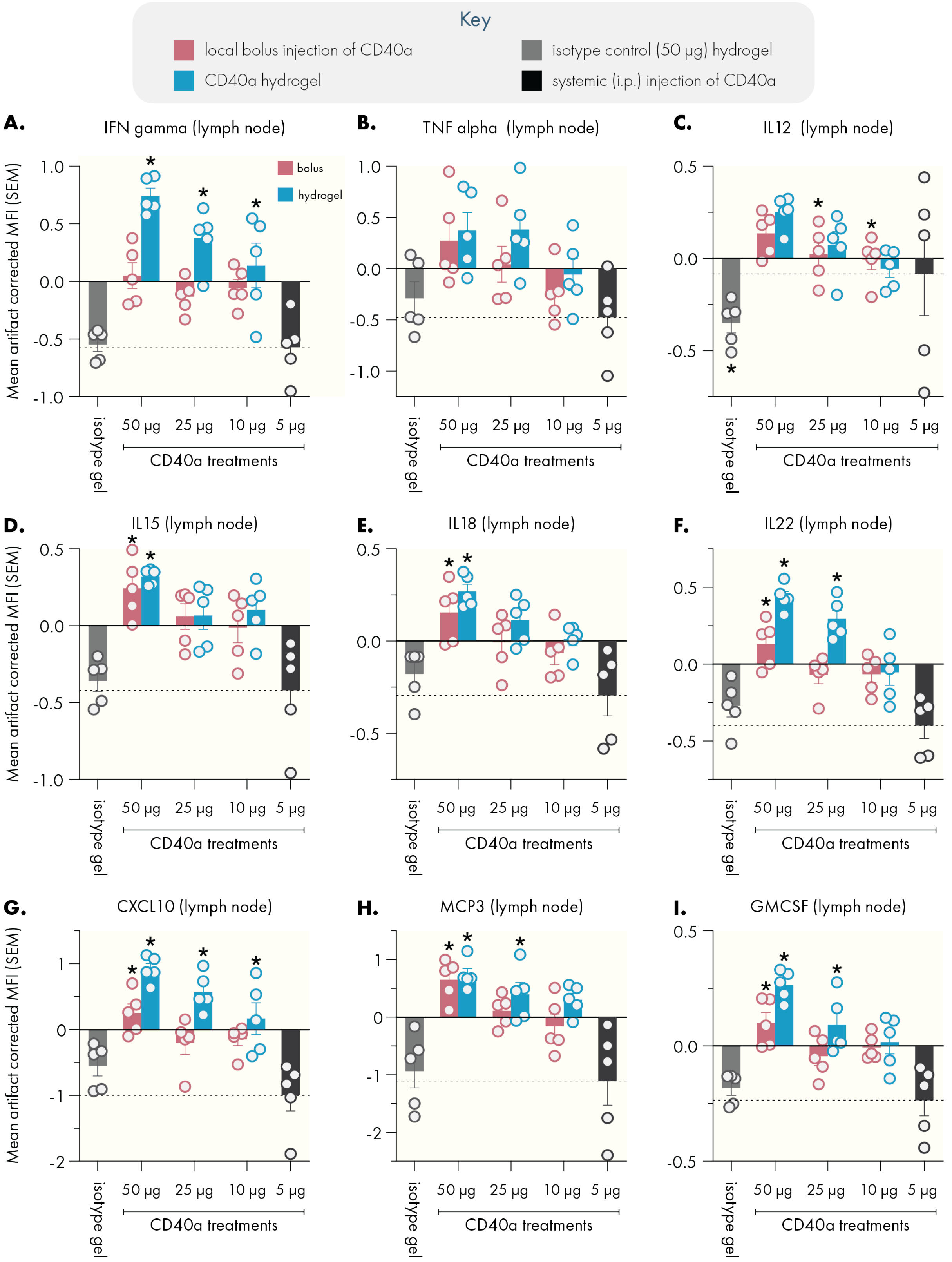


**Figure S19.** Highlighted artifact-corrected lymph node cytokine data. Dotted line indicates mean value corresponding to the 5 μg systemic dose (maximum tolerated systemic dose). N = 5 for all groups, data represented as mean and SEM. * Denote a significant difference ($p < 0.05$) from the MTD systemic dose. Multiple testing error was controlled using the FDR approach (Q = 5%).

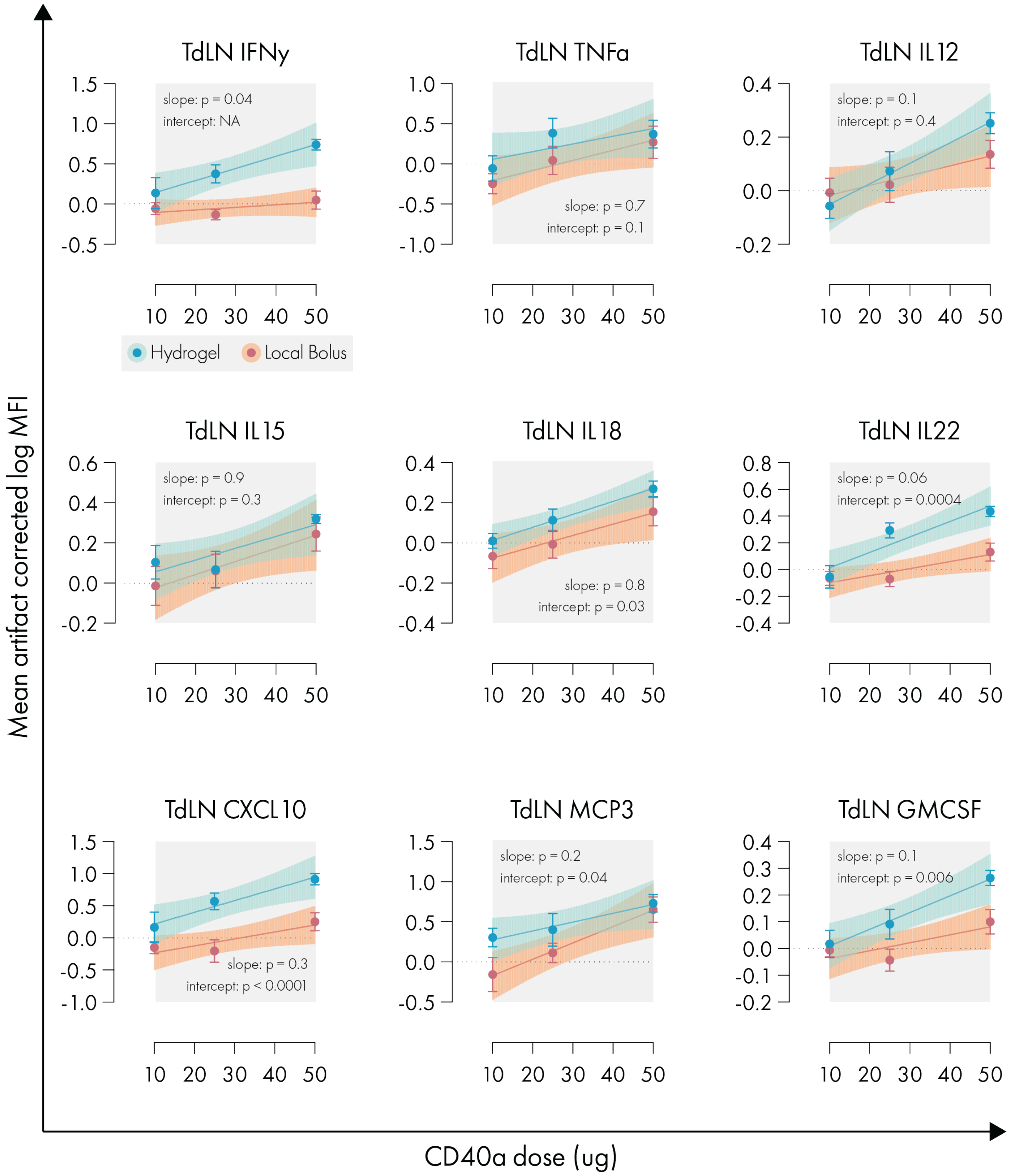


**Figure S20.** Simple linear regression, with no constraints, of dose-TdLN cytokine response curves for individual cytokines. Shaded area represents 95% confidence interval, error bars indicate SEM. Statistical comparisons performed using built-in analysis in the linear regression functionality of GraphPad Prism.

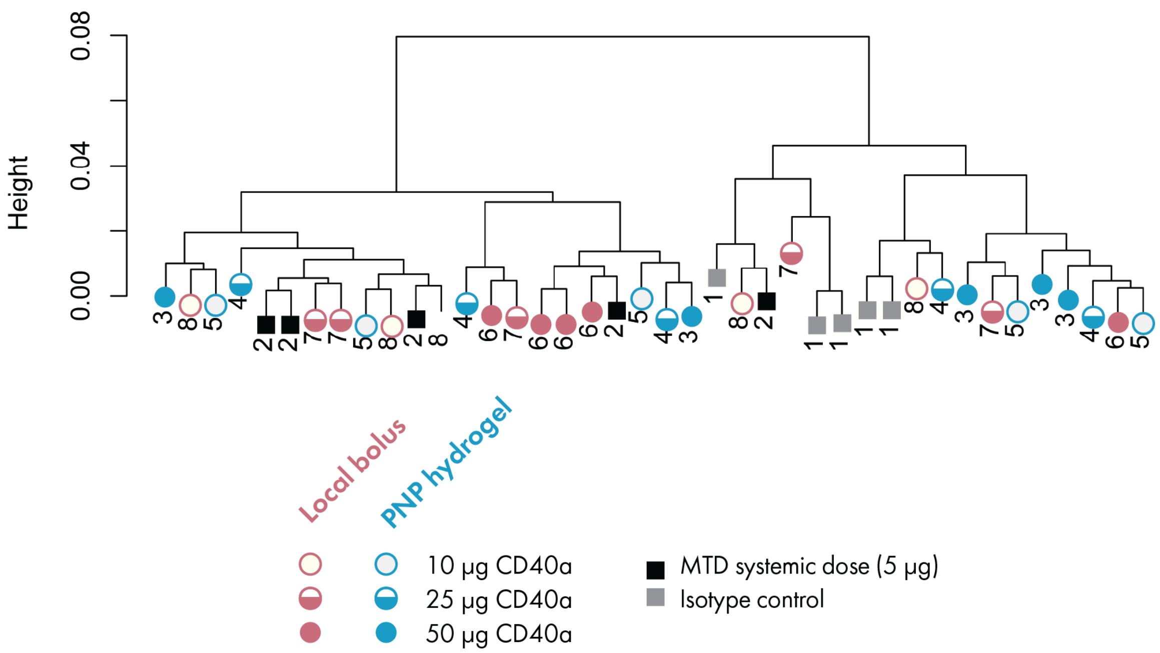


**Figure S21.** TdLN cytokine response dendrogram.
